# Dual function of perivascular fibroblasts in vascular stabilization in zebrafish

**DOI:** 10.1101/2020.04.27.063792

**Authors:** Arsheen M. Rajan, Roger C. Ma, Katrinka M. Kocha, Dan J. Zhang, Peng Huang

## Abstract

Blood vessels are vital to sustain life in all vertebrates. While it is known that mural cells (pericytes and smooth muscle cells) regulate vascular integrity, the contribution of other cell types to vascular stabilization has been largely unexplored. Using zebrafish, we identified sclerotome-derived perivascular fibroblasts as a novel population of blood vessel associated cells. In contrast to pericytes, perivascular fibroblasts emerge early during development, express the extracellular matrix (ECM) genes *col1a2* and *col5a1*, and display distinct morphology and distribution. Time-lapse imaging reveals that perivascular fibroblasts serve as pericyte precursors. Genetic ablation of perivascular fibroblasts results in dysmorphic blood vessels with variable diameters. Strikingly, *col5a1* mutants show spontaneous hemorrhage, and the penetrance of the phenotype is strongly enhanced by the additional loss of *col1a2*. Together, our work reveals dual roles of perivascular fibroblasts in vascular stabilization where they establish the ECM around nascent vessels and function as pericyte progenitors.

**AUTHOR SUMMARY:** Blood vessels are essential to sustain life in humans. Defects in blood vessels can lead to serious diseases, such as hemorrhage, tissue ischemia, and stroke. However, how blood vessel stability is maintained by surrounding support cells is still poorly understood. Using the zebrafish model, we identify a new population of blood vessel associated cells termed perivascular fibroblasts, which originate from the sclerotome, an embryonic structure that is previously known to generate the skeleton of the animal. Perivascular fibroblasts are distinct from pericytes, a known population of blood vessel support cells. They become associated with blood vessels much earlier than pericytes and express several collagen genes, encoding main components of the extracellular matrix. Loss of perivascular fibroblasts or mutations in collagen genes result in fragile blood vessels prone to damage. Using cell tracing in live animals, we find that a subset of perivascular fibroblasts can differentiate into pericytes. Together, our work shows that perivascular fibroblasts play two important roles in maintaining blood vessel integrity. Perivascular fibroblasts secrete collagens to stabilize newly formed blood vessels and a sub-population of these cells also functions as precursors to generate pericytes to provide additional vascular support.

## INTRODUCTION

The vascular system is crucial to the survival of vertebrates. Blood vessels must rapidly expand and contract in response to systemic cues, but also maintain the stability to withstand the stress of blood flow. To maintain their integrity, blood vessels are supported by a highly specialized perivascular architecture comprised of blood vessel associated cells and the surrounding extracellular matrix (ECM) (1,2). Compromised vascular integrity can result in devastating human diseases, such as aneurysms, vascular malformations, and hemorrhagic strokes (3–6). However, how blood vessels are stabilized by different perivascular components is still poorly understood.

The prevailing model is that vascular stability is maintained at three different levels: endothelial cells, mural cells and the surrounding ECM (2). First, blood vessels are lined by endothelial cells. Adherens and tight junctions between endothelial cells provide the primary barrier to passage of fluids, cells, and macromolecules between blood and tissues (7). Second, vascular smooth muscle cells (vSMCs) and pericytes, collectively known as mural cells, closely interact with and stabilize the endothelium (8). vSMCs form a continuous protective sheath around large diameter blood vessels, whereas pericytes are solitary cells partially covering smaller blood vessels such as capillaries. Defects in mural cell specification or recruitment result in widespread vascular leakage and early lethality in mice (9–13). Lastly, the vascular ECM provides structural support for blood vessels (1,14). Mutations in ECM proteins, such as collagens and laminins, lead to early death due to blood vessel rupture in mice (15–18). In humans, defects in fibril forming collagens (type I, III and V), as well as molecules involved in collagen modification and processing, have been associated with Ehlers-Danlos syndrome (EDS) (19). EDS patients are characterized by a significant vascular fragility that often leads to spontaneous rupture of blood vessel walls. Thus, work in both animal models and human patients shows the importance of collagen in maintaining vascular stability. However, one unresolved question is how newly formed blood vessels are stabilized in the early time window before the differentiation of mural cells.

vSMCs and pericytes are the most well-studied blood vessel associated cells and are classically defined by the expression of alpha smooth muscle actin (ACTA2) and platelet-derived growth factor receptor b (PDGFRb), respectively (20,21). Lineage tracing in mice and chick-quail chimeras demonstrate a heterogeneous developmental origin of mural cells (20–23). For example, vSMCs covering the trunk aorta originate from either the neural crest or the sclerotome of the somite, depending on the axial position. Other than mural cells, several types of perivascular cells have been identified, including adventitial cells (24), fibroblasts (25), immune cells (26–28), and astrocytes (29). However, whether these different cell types play roles in vascular stabilization is unknown.

Recent single cell transcriptional profiling studies in mice have revealed the presence of perivascular fibroblast-like cells in the adult mouse brain that are not labeled by classical mural cell markers (30–33). In contrast to mural cells, perivascular fibroblast-like cells adhere loosely to blood vessels and show robust expression of many ECM components, such as collagens. Previous studies have described similar cell populations in the mouse central nervous system that contribute to fibrotic scar formation after injury (34–37). However, where these perivascular fibroblast-like cells originate from during development and how they regulate blood vessel development and stabilization have not been explored.

Zebrafish is a powerful model to study human cardiovascular diseases (38–41). The organization, development and function of the vasculature, including mural cells, are highly conserved between zebrafish and mammals (38,42,43). In the zebrafish trunk, intersegmental vessels (ISVs) sprout from the dorsal aorta (DA) at around 24 hpf (hours post fertilization) and fully establish the metameric pattern by 36 hpf (44). Interestingly, pericytes, the primary mural cells along the ISVs, only emerge at 60 hpf (45). This raises the question how nascent blood vessels are stabilized prior to the differentiation of pericytes. In our work, we describe a novel population of collagen-expressing perivascular fibroblasts that become closely associated with ISVs soon after vessel formation. Perivascular fibroblasts originate from the sclerotome and are distinct from pericytes in their morphology, distribution, and marker expression. Using a combination of *in vivo* live imaging, genetic ablation and CRISPR mutant analysis, we demonstrate that perivascular fibroblasts play dual roles in stabilizing nascent ISVs by regulating the vascular ECM and later functioning as pericyte progenitors. Together, our work provides important insights into the development of perivascular support structures and suggests a molecular and cellular basis of Ehlers-Danlos syndrome.

## RESULTS

### Perivascular fibroblasts express several collagen genes

We previously developed a sclerotome-specific transgenic line *nkx3.1:Gal4; UAS:Nitroreductase-mCherry (nkx3.1^NTR-mCherry^*, similar designations are used for all Gal4/UAS transgenic lines in this paper) (46). This reporter labels the initial sclerotome domains as well as their descendants due to the perdurance of the mCherry protein. Examination of the *nkx3.1^NTR-mCherry^* line in combination with an endothelial reporter, *kdrl:EGFP*, revealed that a population of mCherry^+^ cells was closely associated with intersegmental vessels (ISVs) at 48 hpf (Fig. 1A). This result suggests that the sclerotome contributes to a population of perivascular cells well before the appearance of pericytes at 60 hpf (45). Interestingly, *nkx3.1^NTR-mCherry^*-expressing perivascular cells at 52 hpf were also labeled by a panfibroblast reporter, *col1a2:GFP* (46) (Fig. 1B). Consistent with this result, fluorescent in situ hybridization showed that ISV-associated perivascular cells co-expressed several ECM genes, including *col1a2 (collagen 1a2*) and *col5a1 (collagen 5a1*) at 48 hpf (Fig. 1C and S1). These collagen-expressing perivascular cells are reminiscent of perivascular fibroblast-like cells recently identified in the adult mouse brain by single cell RNA sequencing (Marques et al., 2016; Saunders et al., 2018; Vanlandewijck et al., 2018; Zeisel et al., 2018). We therefore refer to this population of collagen-expressing ISV-associated cells as ‘perivascular fibroblasts’.

**Fig. 1.**
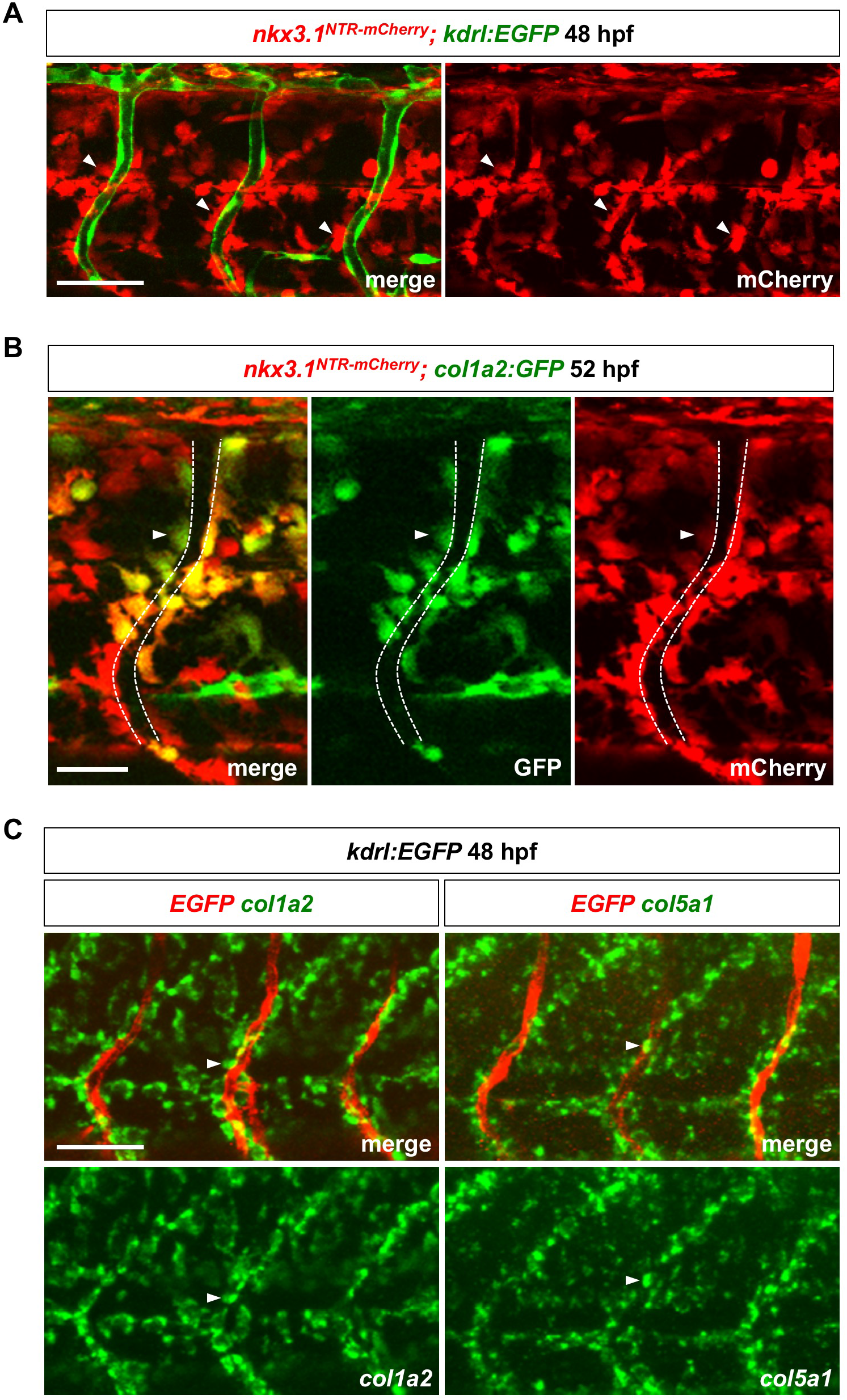
Characterization of perivascular fibroblasts in zebrafish. (A) Lateral view of a three somite region in *nkx3.1^NTR-mCherry^; kdrl:EGFP* embryos at 48 hpf. Many mCherry^+^ perivascular fibroblasts (red, arrowheads) were closely associated with intersegmental vessels (ISVs) labeled by endothelial marker *kdrl:EGFP* (green). *n* = 11 embryos. (B) Co-expression of *nkx3.1^NTR-mCherry^* and *col1a2:GFP* in perivascular fibroblasts (arrowheads) at 52 hpf. The ISV is indicated by dashed lines. *n* = 17 embryos. (C) Fluorescent mRNA in situ hybridization showing expression of fibrillar collagens *col1a2* (green, left) and *col5a1* (green, right) in perivascular fibroblasts (arrowheads) along ISVs marked by *kdrl:EGFP* (red) at 48 hpf. *n* = 15 embryos per staining. Scale bars: (A,C) 50 μm; (B) 25 μm.

### Perivascular fibroblasts originate from the sclerotome

The expression of *nkx3.1^NTR-mCherry^* in perivascular fibroblasts suggests that the sclerotome is the embryonic source of these cells. We have previously shown that the sclerotome in zebrafish has a bipartite organization with a ventral domain and a smaller dorsal domain (46). The ventral sclerotome domain further gives rise to a population of notochord associated cells. At 24 hpf, the sclerotome consists of three compartments: the dorsal sclerotome domain, the ventral sclerotome domain, and notochord associated cells derived from the ventral domain (Fig. 2A). To determine the contribution of these three compartments to perivascular fibroblasts, we performed confocal time-lapse imaging in *nkx3.1^NTR-mCherry^; kdrl:EGFP* embryos. The *nkx3.1^NTR-mCherry^* reporter labels sclerotome progenitors and their progeny, while the endothelial specific *kdrl:EGFP* line allows us to visualize the ISVs. The trunk region of embryos (somite 12-18) was imaged laterally at 6-9 minute intervals starting at 25 hpf, immediately after the emergence of ISV sprouts (Fig. 2B and Movie S1). At 30 hpf, most ISVs became visibly lumenized. During ISV sprouting and lumenization, mCherry^+^ cells can be observed associated with the *kdrl:EGFP-labeled* endothelium (Fig. 2B and S2; Movie S1). By the end of the movie at 49.5 hpf, mCherry^+^ perivascular fibroblasts ‘decorated’ the entire ISVs along the dorsal-ventral (D-V) axis.

**Fig. 2.**
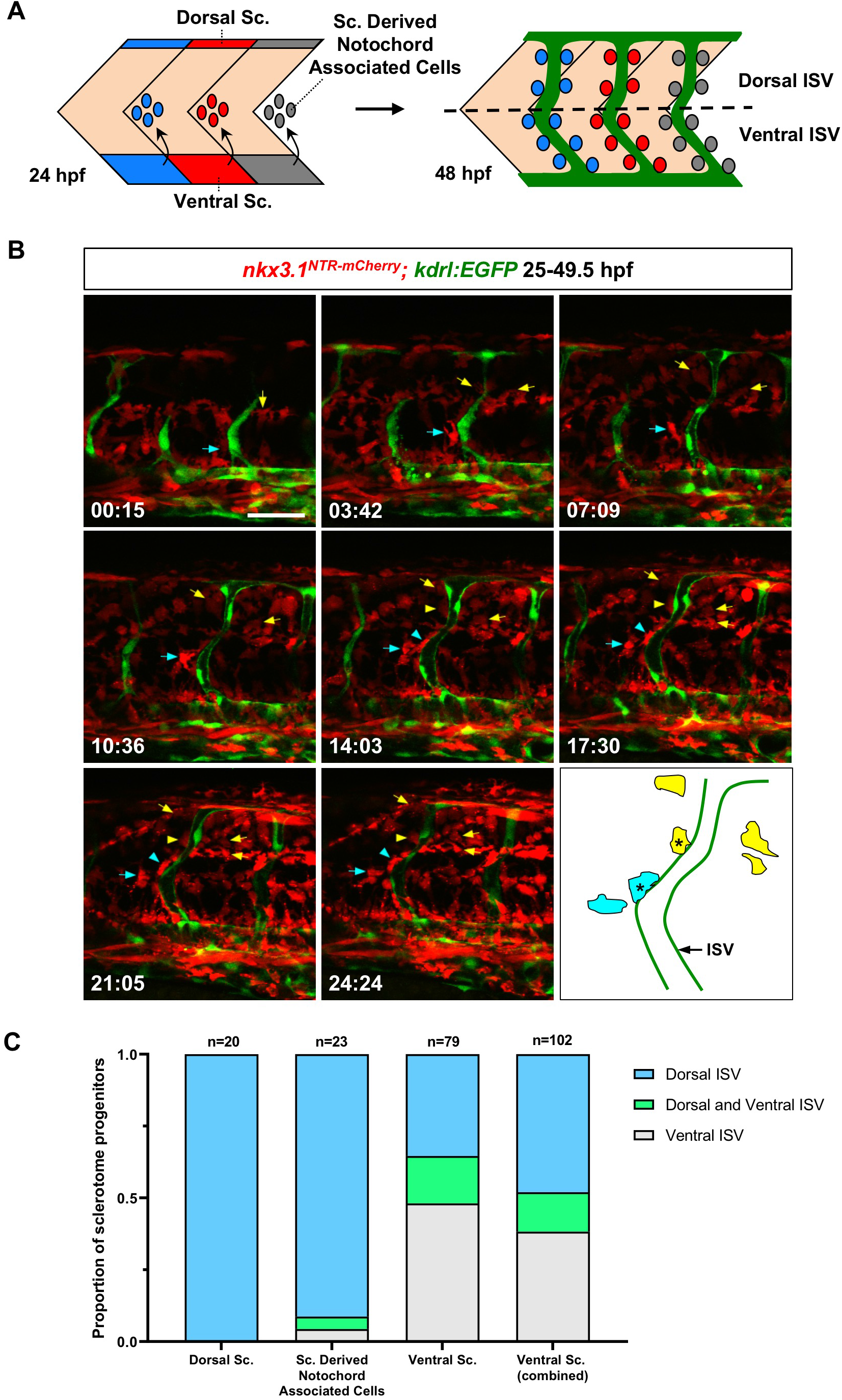
Generation of perivascular fibroblasts from different sclerotome domains. (A) Schematic representation of the bipartite organization of the zebrafish sclerotome and the generation of perivascular fibroblasts. At 24 hpf, the zebrafish sclerotome in each somite is divided into three compartments: the dorsal sclerotome, the ventral sclerotome, and sclerotome derived notochord associated cells. Note that notochord associated cells originate from the ventral sclerotome and are located about half-somite posterior to the corresponding somite. At 48 hpf, perivascular fibroblasts appear along the length of the intersegmental vessels (ISVs, green). The dotted line indicates the position of the horizontal myoseptum, which divides any given ISV into a dorsal half and a ventral half. Perivascular fibroblasts were quantified based on their final locations along each ISV: dorsal ISV (above the horizontal myoseptum) or ventral ISV (below the horizontal myoseptum). (B) Snapshots from time-lapse imaging of a *nkx3.1^NTR-mCherry^; kdrl:EGFP* embryo from 25 hpf to 49.5 hpf. Perivascular fibroblasts along ISVs were retrospectively traced to determine their cell of origin. One cell from the ventral sclerotome domain (cyan arrows) and one cell from sclerotome derived notochord associated cells (yellow arrows) were traced over 24.5 hours with their daughter cells indicated by the same colored arrows/arrowheads. Both sclerotome progenitors divided at least once to give rise to one perivascular fibroblast (arrowheads) as well as several interstitial cells (arrows). A schematic representation of color-coded traced cells at the last time point is shown with perivascular fibroblasts indicated by asterisks. The corresponding time-lapse movie is shown in Movie S1. *n* = 6 embryos. (C) Quantification of the contribution of each sclerotome domain to perivascular fibroblasts. Sclerotome progenitors from each domain were quantified based on the final dorsoventral location of perivascular fibroblasts along each ISV. A given sclerotome progenitor can give rise to perivascular fibroblasts in only dorsal ISV, in only ventral ISV, or in both dorsal and ventral ISV as indicated in (A). The ventral sclerotome (combined) group includes progenitors from both the ventral sclerotome domain and sclerotome derived notochord associated cells. *n* = 122 sclerotome progenitors from 6 embryos. Scale bar: 50 μm.

By retrospective cell tracing, we showed that perivascular fibroblasts can be traced to all three compartments of the sclerotome. Of 170 perivascular fibroblasts traced, 28 cells originated from the dorsal sclerotome domain (16%), 112 from the ventral sclerotome domain (66%), and 30 from notochord associated cells (18%). Since notochord associated cells are derived from the ventral sclerotome domain, this result suggests that the ventral sclerotome is the main contributor of perivascular fibroblasts (142/170 cells, 84%).

To better compare the contribution of sclerotome progenitors from different compartments, we subdivided perivascular fibroblasts into two domains based on their final D-V positions along the ISVs. The position of the horizontal myoseptum was used as a landmark to define perivascular fibroblasts at dorsal or ventral ISVs (Fig. 2A). The 170 perivascular fibroblasts can be traced back to 122 sclerotome progenitors. These sclerotome progenitors typically underwent 1-2 cell divisions over the 24-hour period to give rise to multiple daughter cells, at least one of which became associated with the neighboring ISV. We categorized sclerotome progenitors into three groups based on the final positions of perivascular fibroblasts they generated: 1) only at the dorsal ISV, 2) only at the ventral ISV, or 3) at both dorsal and ventral ISV (Fig. 2C). Sclerotome progenitors in the dorsal domain gave rise to exclusively perivascular fibroblasts along dorsal ISVs (20/20 cells, 100%) (Fig. 2C). Similarly, the majority of sclerotome progenitors around the notochord (i.e., sclerotome derived notochord associated cells) (21/23 cells, 91%) contributed to only perivascular fibroblasts at the dorsal ISV (Fig. 2C). By contrast, 48% of sclerotome progenitors in the ventral domain (38/79 cells) gave rise to only perivascular fibroblasts at the ventral ISV, 35% of ventral sclerotome progenitors (28/79 cells) generated only dorsal perivascular fibroblasts, while the remaining 16% of ventral progenitors (13/79 cells) contributed to perivascular fibroblasts at both dorsal and ventral ISV regions (Fig. 2C). Similar to the contribution of tendon fibroblasts (tenocytes) (46), the dorsal sclerotome generates only dorsally positioned perivascular fibroblasts, whereas the ventral sclerotome contributes to perivascular fibroblasts along the entire D-V axis of ISVs (the combined bar in Fig. 2C). Moreover, perivascular fibroblasts along a given ISV were always derived from the sclerotome of the same overlying somite (170/170 cells, 100%). Together, our results demonstrate that perivascular fibroblasts originate from all three sclerotome compartments in a stereotypic manner.

### Perivascular fibroblasts are distinct from pericytes

Mural cells, including pericytes and vSMCs, are known blood vessel associated support cells. We next asked whether perivascular fibroblasts represent a different perivascular cell population from mural cells. In the zebrafish trunk, pericytes are associated with ISVs, while vSMCs are localized to larger vessels such as the dorsal aorta (43,45). Platelet derived growth factor (PDGF) signaling is known to play an important role in pericyte recruitment and the platelet derived growth factor receptor-beta *(pdgfrb*) is a well-established pericyte marker (45; 47–48). We utilized the *pdgfrb:Gal4FF; UAS:NTR-mCherry* line (referred to as *pdgfrb^NTR-mCherry^*) to label pericytes, while the *nkx3.1^NTR-mCherry^* and *col1a2:GFP* reporters were used to mark perivascular fibroblasts. Our results indicate that perivascular fibroblasts and pericytes represent distinct cell populations. First, they showed different developmental timing and cell numbers. Consistent with our time-lapse movies, *nkx3.1^NTR-mCherry^-* positive perivascular fibroblasts emerged along ISVs prior to 2 dpf (days post fertilization) (Fig. 1A and S2) and remained largely constant in number (9.9-10.7 per ISV) from 2 to 4 dpf (Fig. 3A). By contrast, pericytes marked by *pdgfrb^NTR-mCherry^* did not appear on ISVs until 3 dpf, and the cell number increased to 0.9 per ISV by 4 dpf (Fig. 3A), similar to what has been previously described (45). Second, perivascular fibroblasts and pericytes showed distinct marker expression (Fig. 3B). Perivascular fibroblasts labeled by *col1a2:GFP* did not express the *pdgfrb^NTR-mCherry^* reporter at 4 dpf. Conversely, *pdgfrb^NTR-mCherry^-positive* pericytes showed minimal *col1a2:GFP* expression. While perivascular fibroblasts appeared globular, pericytes were more elongated with long cellular processes (Fig. 3B). Moreover, perivascular fibroblasts were more laterally located along ISVs compared to pericytes, with pericyte processes often sandwiched between the neighboring perivascular fibroblasts and the endothelium (Fig. 3B). To further examine the morphological differences between perivascular fibroblasts and pericytes, we performed high resolution confocal microscopy at the single cell resolution at 3 dpf. *pdgfrb^NTR-mCherry^-positive* pericytes showed the typical pericyte morphology: small and flat cell bodies with several elongated cellular processes that extended longitudinally along and tightly wrapped around the ISV, labeled by *kdrl:EGFP* (Fig. 3C). Since perivascular fibroblasts are more abundant along ISVs compared to pericytes, we took advantage of a mosaic *col1a2:Gal4; UAS:Kaede* line (referred to as *col1a2^Kaede^*) (49) to sparsely label single perivascular fibroblasts for easier visualization. In contrast to pericytes, perivascular fibroblasts were more loosely associated with ISVs and had a globular morphology with shorter and finer processes that wrapped diametrically around the associated ISV in an ‘awkward hug’ (Fig. 3D). Taken together, our results suggest that perivascular fibroblasts and pericytes represent two distinct populations of perivascular cells with unique marker expression, morphology, distribution and developmental timing.

**Fig. 3.**
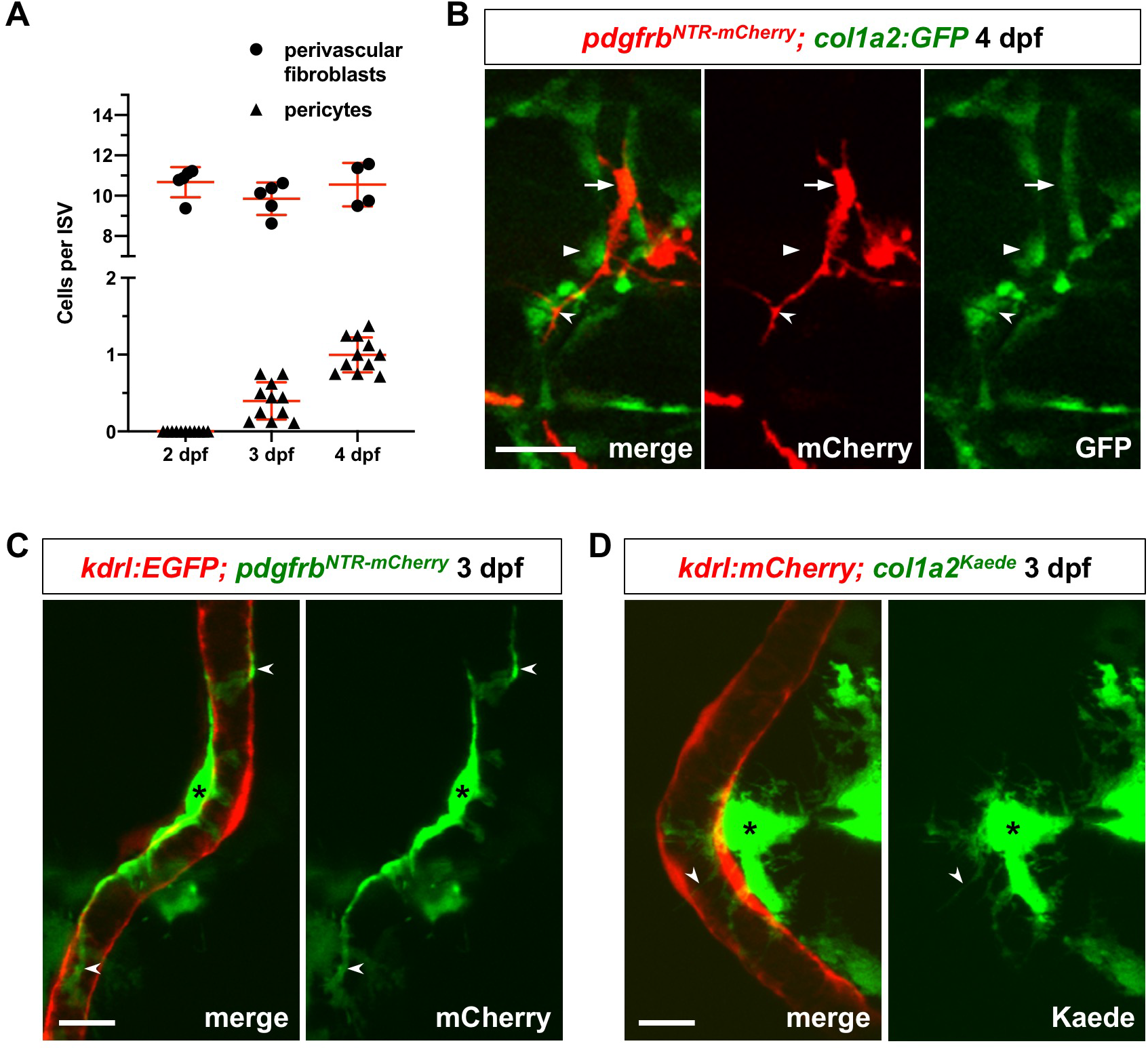
Perivascular fibroblasts are distinct from pericytes. (A) Quantification of the number of perivascular fibroblasts and pericytes during development. The number of perivascular fibroblasts and pericytes were scored in *nkx3.1^NTR-mCherry^; kdrl:EGFP* and *pdgfrb^NTR-mCherry^; kdrl:EGFP* embryos, respectively. Each data point represents the average cell number of 8-10 ISVs from an individual embryo. Data are plotted with mean ± SEM indicated. *n* = 4-5 (*nkx3.1^NTR-mCherry^; kdrl:EGFP*) and 11 (*pdgfrb^NTR-mCherry^; kdrl:EGFP*) embryos at each time point. (B) *pdgfrb^NTR-mCherry^; col1a2:GFP* embryos were imaged at 4 dpf to visualize perivascular fibroblasts and pericytes. No overlap in marker expression was observed between *pdgfrb^NTR-mCherry^-positive* pericytes (red, arrows) and *col1a2:GFP-* positive perivascular fibroblasts (green, arrowheads). Unlike perivascular fibroblasts, pericytes also displayed elongated cellular processes (notched arrowheads). *n* = 14 embryos. (C) *pdgfrb^NTR-mCherry^; kdrl:EGFP* embryos were imaged at 3 dpf to visualize individual pericytes (green, asterisks) with long cellular processes (notched arrowheads) that wrapped around the ISV (red). (D) Mosaic *col1a2^Kaede^* line was imaged to visualize a single perivascular fibroblast (green, asterisks) associated with an ISV (red) in *col1a2^Kaede^*; *kdrl:mCherry* embryos at 3 dpf. Perivascular fibroblasts showed an overall globular morphology with shorter processes (notched arrowheads) wrapping loosely around the neighbouring ISV. Scale bars: (B) 25 μm; (C,D) 10 μm.

### A sub-population of perivascular fibroblasts functions as pericyte progenitors

While occupying the similar perivascular space, perivascular fibroblasts appear along ISVs at least one day before pericytes (Fig. 3A). We hypothesized that perivascular fibroblasts act as progenitors for ISV pericytes. To test this possibility, we conducted time-lapse imaging of *pdgfrb^NTR-mCherry^; col1a2:GFP* embryos from 54 to 73 hpf during which pericytes started to appear on ISVs.

Interestingly, a subset of *col1a2:GFP*-positive perivascular fibroblasts gradually lost GFP expression, switched on the pericyte reporter *pdgfrb^NTR-mCherry^,* and developed long cellular processes characteristic of pericytes at this stage (Fig. 4A and Movie S2). Indeed, of 15 newly formed pericytes traced in our time-lapse movies, 60% were derived from *col1a2:GFP-expressing* perivascular fibroblasts (9/15 cells). This result suggests that a sub-population of perivascular fibroblasts can differentiate into pericytes. Since *col1a2* transgenic lines such as *col1a2:GFP* are mosaic (Fig. S3A) (49), the number above likely under-estimates the proportion of pericytes derived from perivascular fibroblasts. To further test our model, we performed time-lapse experiments in *nkx3.1^NTR-mCherry^; pdgfrb:GFP* embryos. Interestingly, while both *pdgfrb:GFP* and *pdgfrb^NTR-mCherry^* lines labeled pericytes (GFP^high^mCherry^+^ expression) at 4 dpf, *pdgfrb:GFP* but not *pdgfrb^NTR-mCherry^* also marked perivascular fibroblasts (GFP^low^mCherry^−^ expression) at 2 and 4 dpf (Fig. S3B and S3C). This is consistent with the previous report that the sclerotome expresses *pdgfrb* (50), suggesting that *pdgfrb:GFP* is a more sensitive reporter than the *pdgfrb^NTR-mCherry^* line. Similar to movies in *pdgfrb^NTR-mCherry^; col1a2:GFP* embryos, cell tracing in *nkx3.1^NTR-mCherry^; pdgfrb:GFP* embryos showed that some mCherry^+^GFP^low^ perivascular fibroblasts slowly upregulated GFP expression, and extended pericyte-like cellular processes wrapping around the ISV (Movie S3). Of 117 newly formed GFP^high^ pericytes, 91% of them (106/117 cells) were derived from mCherry^+^ perivascular fibroblasts. This result confirms that a subpopulation of perivascular fibroblasts functions as pericyte precursors. Since perivascular fibroblasts originate from the sclerotome, we conclude that most pericytes along ISVs are also derived from the sclerotome.

**Fig. 4.**
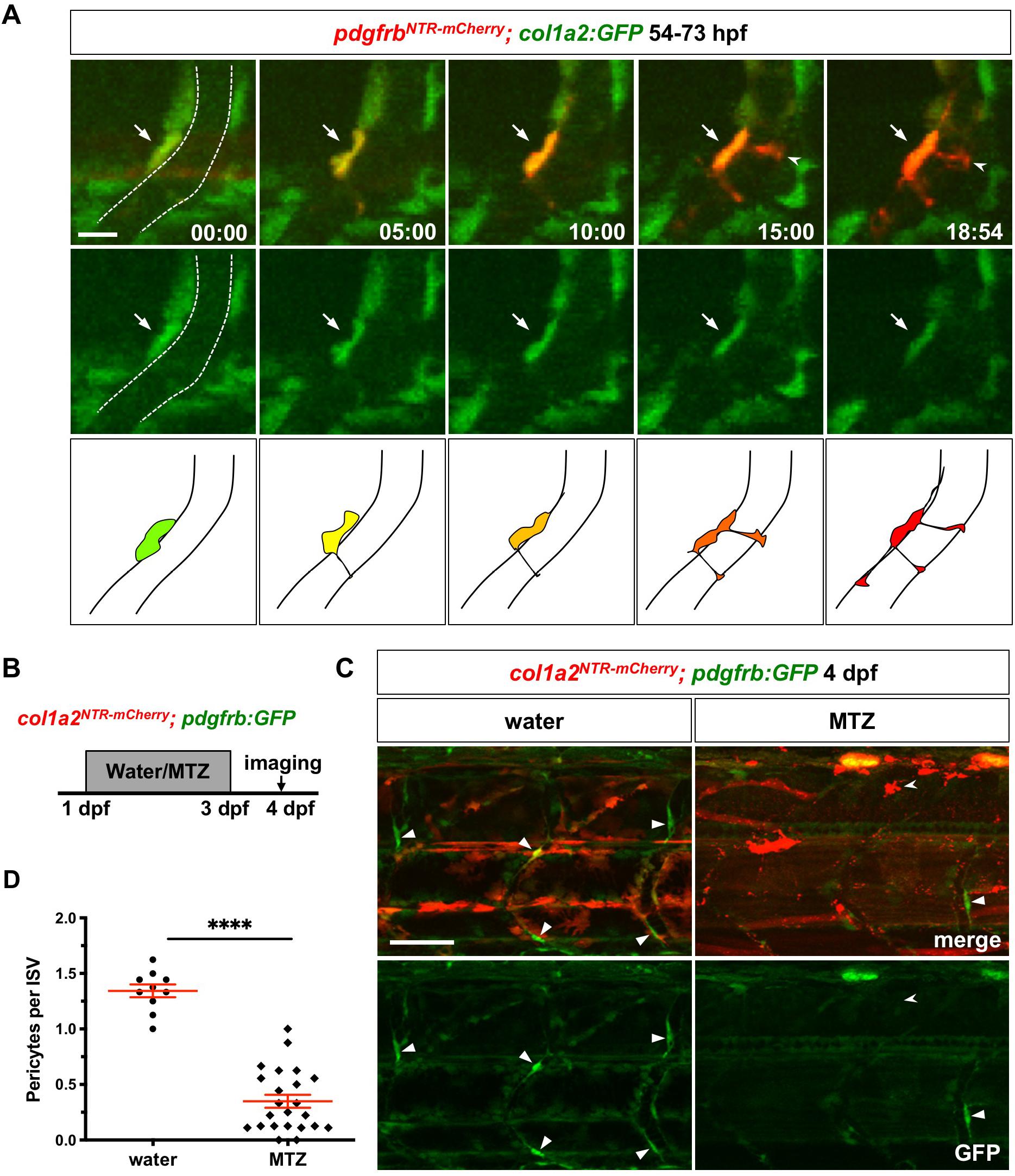
Perivascular fibroblasts function as pericyte progenitors. (A) Snapshots from time-lapse imaging of *pdgfrb^NTR-mCherry^; col1a2:GFP* embryos from 54 to 73 hpf. Newly differentiated pericytes were retrospectively traced to identify their cell of origin. The ISV is outlined by dotted lines at the first time point. One perivascular fibroblast (green, arrows) traced can be seen gradually upregulating *pdgfrb^NTR-mCherry^* expression and extending pericyte-like cellular processes (notched arrowheads). The time stamps are indicated in the hh:mm format. Schematic drawings of the merged images at each time point are shown at the bottom. The corresponding time-lapse movie is shown in Movie S2. *n* = 7 embryos. (B) Schematic showing experimental timeline of perivascular fibroblast ablation. *col1a2^NTR-mCherry^; pdgfrb:GFP* embryos were incubated in either water or metronidazole (MTZ) from 1 to 3 dpf following which embryos was washed at 3 dpf and imaged at 4 dpf to visualize pericytes. (C) Representative images of water (left) or MTZ (right) treated *col1a2^NTR-mCherry^; pdgfrb:GFP* embryos at 4 dpf. Compared to water-treated control embryos, MTZ-treated embryos showed complete ablation of mCherry^+^ cells with only residual mCherry^+^ debris (notched arrowheads). Fewer *pdgfrb:GFP^high^* pericytes (arrowheads) can be seen in MTZ-treated embryos compared to water-treated controls. (D) Quantification of the number of pericytes after perivascular fibroblast ablation. Pericytes were scored based on high level expression of the *pdgfrb:GFP* reporter as shown in (C). Each data point represents the average number of pericytes per ISV scored from 8-10 ISVs from an individual embryo. *n* = 10 (water) and 23 (MTZ) embryos. Data are plotted as mean ± SEM. Statistics: Mann-Whitney *U* test. Asterisk representation: p-value < 0.0001 (****). Scale bars: (A) 10 μm; (C) 50 μm.

To further test if perivascular fibroblasts serve as pericyte precursors, we performed genetic ablation of perivascular fibroblasts using the *col1a2:Gal4; UAS:NTR-mCherry* line (referred to as *col1a2^NTR-mCherry^*). Nitroreductase (NTR) converts metronidazole (MTZ), an otherwise harmless prodrug into a cytotoxic compound, resulting in the death of NTR-expressing cells (51). To ablate perivascular fibroblasts prior to pericyte differentiation, *col1a2^NTR-mCherry^; pdgfrb:GFP* embryos were treated with MTZ from 1 to 3 dpf, after which the drug was washed off and embryos were imaged for pericytes at 4 dpf (Fig. 4B). In contrast to water-treated controls, we observed a complete loss of perivascular fibroblasts in MTZ-treated embryos, with only residual mCherry^+^ debris visible in the trunk (Fig. 4C). Interestingly, we observed a drastic 3.5-fold reduction in the number of pericytes in MTZ-treated embryos (0.4 cells per ISV) compared to water-treated controls (1.4 cells per ISV) (Fig. 4C and 4D). This result is consistent with our model that perivascular fibroblasts function as pericyte precursors as depletion of perivascular fibroblasts leads to the decrease in the number of pericytes. Together, our work suggests that the sclerotome generates perivascular fibroblasts, some of which further differentiate into pericytes along the ISVs.

### Loss of perivascular fibroblasts results in dysmorphic ISVs

Our work suggests that perivascular fibroblasts function as progenitors to generate pericytes to support the vasculature. However, only a small fraction of perivascular fibroblasts actually differentiates into pericytes, raising the question about the function of the remaining perivascular fibroblasts. Moreover, the emergence of perivascular fibroblasts occurs concurrently with ISV development (Fig. 2B and S2; Movie S1), well before pericyte differentiation at 60 hpf (45). These observations raise the possibility that perivascular fibroblasts play an early role in stabilizing nascent blood vessels prior to pericyte formation. To test this idea, we examined the impact of early ablation of perivascular fibroblasts on ISV development using the nitroreductase-based system. *nkx3.1^NTR-mCherry^; kdrl:EGFP* embryos were incubated in water or MTZ for a 24 hour period starting at 38 hpf when ISV formation was complete and blood flow had commenced in ISVs (Fig. 5A). ISV morphology was examined after the drug treatment at 62 hpf. Water-treated control embryos showed stereotypical ISVs with many mCherry^+^ perivascular fibroblasts (Fig. 5B). MTZ treatment resulted in a complete loss of perivascular fibroblasts, with only some mCherry^+^ cell debris remaining (Fig. 5B). Compared to control ISVs, most ISVs in ablated embryos showed visible distortions in their diameters, shrinking and dilating dramatically at different points along the ISV length (Fig. 5B). To quantify this variability, the diameter of each ISV was measured at four equidistant points along its length and standard deviation from the mean diameter was graphed as a readout for ISV diameter variability (Fig. 5C). Indeed, ISVs in the absence of perivascular fibroblasts were significantly more variable than those of water-treated controls (Fig. 5C). Interestingly, similar ablation experiments using *col1a2^NTR-mCherry^; kdrl:EGFP* during later stages between 4 and 5 dpf did not significantly alter the ISV morphology (Fig. S4). Together, our results suggest that perivascular fibroblasts play an early role in the stabilization of nascent blood vessels but are dispensable for maintaining ISV morphology at later stages.

**Fig. 5.**
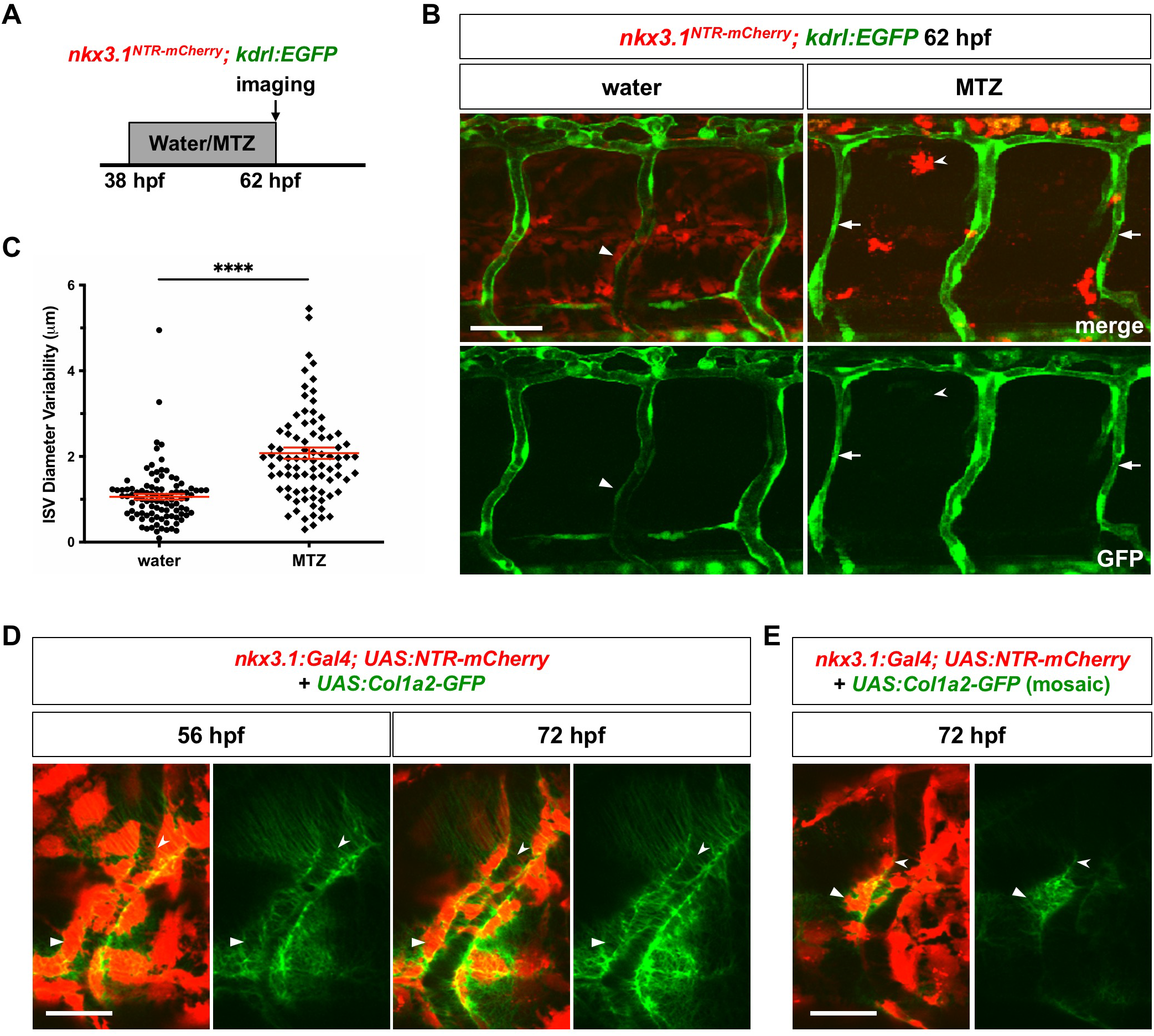
Perivascular fibroblasts stabilize nascent blood vessels by collagen deposition. (A) Schematic of experimental protocol for early ablation of perivascular fibroblasts. *nkx3.1^NTR-mCherry^; kdrl:EGFP* embryos were incubated in either water or metronidazole (MTZ) from 38 to 62 hpf and imaged to visualize ISV morphology. (B) Representative images showing water (left) and MTZ (right) treated embryos. Water-treated control embryos had many mCherry^+^ cells (arrowheads), while MTZ treatment resulted in complete ablation of mCherry^+^ cells with only mCherry^+^ debris (notched arrowheads) remaining. MTZ-treated embryos showed visibly deformed ISV morphology with greater variation in vessel diameter (arrows indicate the narrow region of ISVs) compared to uniform ISVs in controls. (C) Quantification of ISV diameter variability in (B). ISV diameter was measured at 4 equidistant points along each ISV using the line tool in ImageJ. Mean diameter of each ISV was calculated and standard deviation from the mean was plotted as a readout of diameter variability in each ISV examined. MTZ-treated embryos showed significantly more variable ISVs compared to water-treated controls. *n* = 115 ISVs from 14 embryos (water); 126 ISVs from 15 embryos (MTZ). Results were graphed as mean ± SEM. Statistics: Mann-Whitney *U* test. Asterisk representation: p-value < 0.0001 (****). (D) *nkx3.1^NTR-mCherry^* embryo were injected with the *UAS:Col1a2-GFP* plasmid and imaged at 56 and 72 hpf. Many mCherry^+^GFP^+^ perivascular fibroblasts (arrowheads) can be seen surrounding the ISV. Numerous thin GFP^+^ collagen fibers (notched arrowheads) wrapped around the ISV and the Col1a2-GFP protein deposition appeared to increase from 56 to 72 hpf. *n* = 16 embryos. (E) *nkx3.1^NTR-mCherry^* embryos were injected with a low dose of the *UAS:Col1a2-GFP* plasmid and imaged at 72 hpf. Col1a2-GFP deposition (notched arrowheads) around an ISV by a single mCherry^+^GFP^+^ perivascular fibroblast (arrowheads) can be seen. *n* = 16 embryos. Scale bars: (B) 50 μm; (D,E) 25 μm.

### Perivascular fibroblasts deposit collagens around ISVs

Our work suggests that perivascular fibroblasts are crucial for the stabilization of nascent ISVs. Since perivascular fibroblasts show high-level expression of collagen genes, including *col1a2* and *col5a1,* we predicted that perivascular fibroblasts are the main source of the vascular ECM around newly formed ISVs. To test this model, we generated a *UAS:Col1a2-GFP* construct to express GFP-fused Col1a2 in live embryos. A similar *Col1a2-GFP* reporter has been previously used to visualize collagen deposition during skin development and repair in zebrafish (52). *nkx3.1:Gal4; UAS:NTR-mCherry* embryos injected with the *UAS:Col1a2-GFP* plasmid showed many mCherry^+^GFP^+^ perivascular fibroblasts surrounding ISVs at 56 hpf (Fig. 5D). Indeed, numerous thin GFP^+^ ‘strings’, likely corresponding to collagen fibers, wrapped around the ISV with the intensity increasing from 56 to 72 hpf, suggesting continuous Col1a2-GFP protein deposition (Fig. 5D). To determine whether perivascular fibroblasts directly contribute to collagens around ISVs, we injected *nkx3.1^NTR-mCherry^* embryos with a low dose of the *UAS:Col1a2-GFP* plasmid to achieve mosaic labeling of perivascular fibroblasts. Imaging of individually labeled mCherry^+^GFP^+^ perivascular fibroblasts showed local Col1a2-GFP deposition around the ISV within 1-2 cell diameters from the Col1a2-GFP-expressing perivascular fibroblast (Fig. 5E). Our results suggest that perivascular fibroblasts contribute to collagens in the ECM surrounding nascent ISVs that likely contribute to vessel stabilization.

### *collagen* mutants display severe ISV hemorrhage

Our work suggests that perivascular fibroblasts secrete collagens and stabilize nascent ISVs. We next asked whether Col1a2 and Col5a1, which are expressed in perivascular fibroblasts (Fig. 1C and S1), are required to maintain the integrity of nascent blood vessels. Using CRISPR/Cas9 editing (53), we generated mutants for *col1a2* (*col1a2^ca108^*, 1-bp deletion) and *col5a1* (*col5a1^ca109^*, 4-bp deletion) (Fig. S5A and S5B). Both alleles resulted in frame shifts in the corresponding coding sequences leading to premature stops (Fig. S5C and S5D). For simplicity, we designate wild-type, heterozygous or homozygous fish as *+/+, +/−*, and *−/−*, respectively. For example, heterozygous carriers of *col1a2^ca108^* are referred to as *col1a2^+/−^.* Whole mount in situ hybridization showed that mutant mRNAs for both genes underwent nonsense-mediated decay (Fig. 6A), suggesting that both *col1a2^ca108^* and *col5a1^ca109^* are null alleles. Crosses of heterozygous *col1a2^+/−^* parents gave rise to progeny segregated in roughly Mendelian ratios at all stages examined (Fig. S6A). Although homozygous *col1a2^−/−^* mutants were viable as adults, they were substantially smaller than their wild-type siblings and often showed severe spine curvature (Fig. S6B). This phenotype is reminiscent of those described in other type I collagen mutants (54,55). By contrast, from crosses of heterozygous *col5a1^+/−^* parents, homozygous *col5a1^−/−^* mutants were under-represented at 11 dpf, and completely absent by 14 dpf (Fig. S6C). These results suggest that Col5a1 is a more critical component of the ECM compared to Col1a2, and it is required for the survival of the fish to adulthood. Strikingly, a small percentage of homozygous *col5a1^−/−^* embryos showed spontaneous hemorrhage in the trunk region at 2 dpf, suggesting compromised vascular integrity in the absence of Col5a1 (Fig. 6B).

**Fig. 6.**
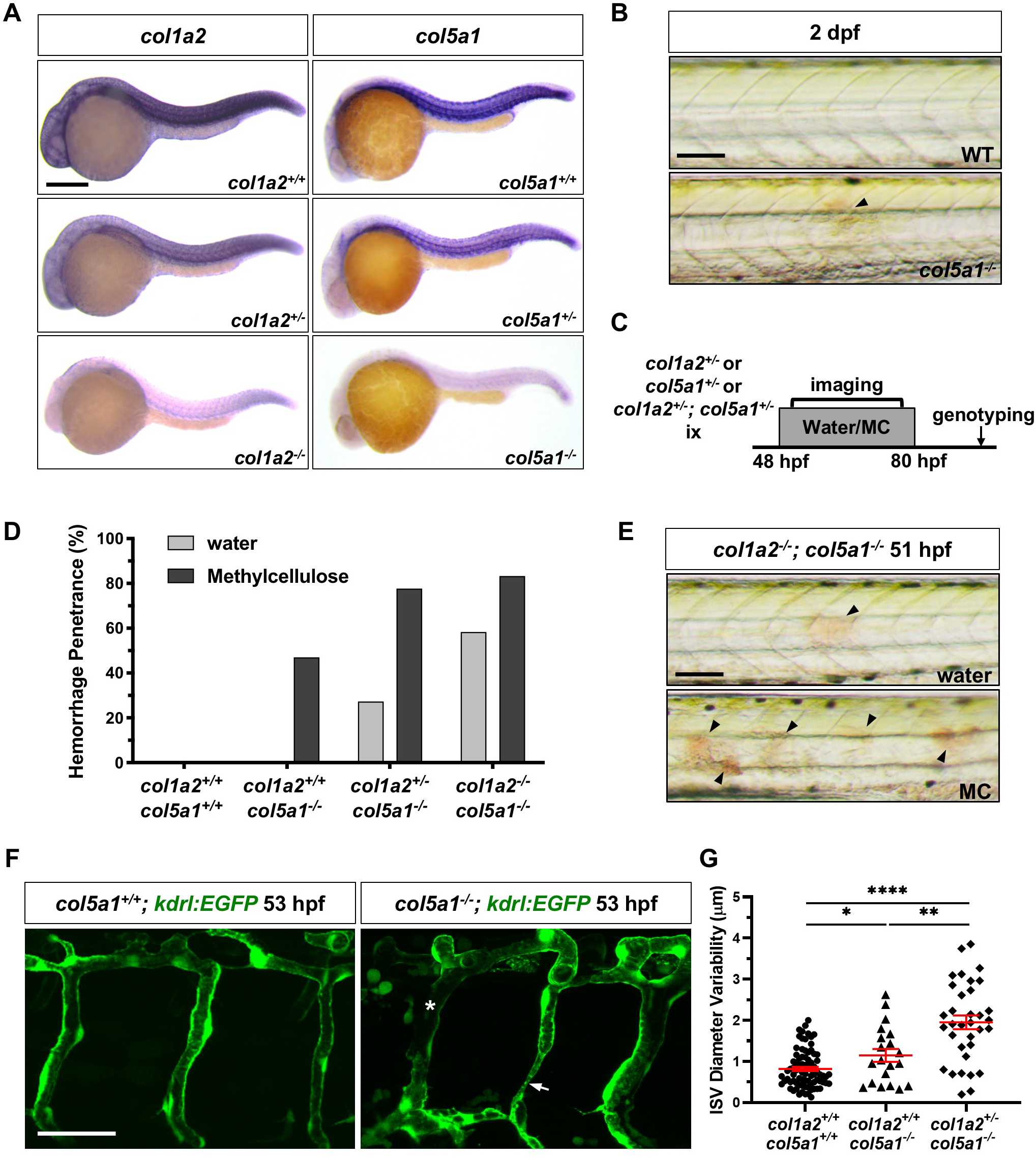
Characterization of collagen mutants. (A) Embryos from intercrosses of *col1a2^+/−^* adults or *col5a1^+/−^* adults were stained at 24 hpf by mRNA in situ hybridization with *col1a2* (left) or *col5a1* (right) probes, respectively. Compared to wild type siblings, heterozygous mutants showed reduced staining, and homozygous mutants displayed an almost complete loss of staining for both genes examined. *n* = 30 embryos for each staining. (B) *col5a1^−/−^* mutants but not wild-type siblings showed spontaneous hemorrhage in the trunk (arrowhead) at 2 dpf. (C) Schematic of experimental protocol for phenotypic analysis of collagen mutants. Embryos from intercrosses of 1) *col1a2^+/−^* adults, 2) *col5a1^+/−^* adults, or 3) *col1a2^+/−^; col5a1^+/−^* adults were incubated in either water or 0.6% methylcellulose (MC) and screened for the hemorrhage phenotype in the trunk from 48 to 80 hpf. All embryos were subsequently genotyped. (D) Quantification of hemorrhage penetrance of embryos from intercrosses of *col1a2^+/−^; col5a1^+/−^* adults described in (C). The hemorrhage penetrance was calculated by dividing the number of embryos with the hemorrhage phenotype by the total number of embryos of the same genotype. *n* = 12 *(col1a2^+/+^; col5a1^+/+^* + water); 9 *(col1a2^+/+^; col5a1^+/+^* + MC); 12 *(col1a2^+/+^; col5a1^−/−^* + water); 12 *(col1a2^+/+^; col5a1^−/−^* + MC); 22 *(col1a2^+/−^; col5a1^−/−^* + water); 27 *(col1a2^+/−^; col5a1^−/−^* + MC); 12 *(col1a2^−/−^; col5a1^−/−^* + water); and 18 *(col1a2^−/−^; col5a1^−/−^* + MC) embryos. (E) *col1a2^−/−^; col5a1^−/−^*embryos were incubated in water (top) or MC (bottom) for 3 hours at 2 dpf. MC-treated *col1a2^−/−^; col5a1^−/−^* embryos showed an increase in the number of hemorrhage foci (arrowheads) compared to water-treated controls. Quantification of this result is shown in Fig S6D. (F) Embryos from crosses of *col1a2^+/−^; col5a1^+/−^* and *col5a1^+/−^; kdrl:EGFP* adults were incubated in the 0.6% methylcellulose solution at 48 hpf, and their ISVs were imaged at 53 hpf. *col5a1^−/−^; kdrl:EGFP* embryos showed visible ISV constrictions (arrow) and broken ISVs (asterisk) compared to *col5a1^+/+^; kdrl:EGFP* siblings. (G) Quantification of ISV diameter variability in embryos from crosses of *col1a2^+/−^; col5a1^+/−^* and *col5a1^+/−^; kdrl:EGFP* adults as described in (F). ISV diameter and variability were measured as described in Fig 5. *n* = 81 ISVs from 10 embryos *(col1a2^+/+^; col5a1^+/+^; kdrl:EGFP);* 20 ISVs from 3 embryos *(col1a2^+/+^; col5a1^−/−^; kdrl:EGFP);* and 34 ISVs from 4 embryos *(col1a2^+/−^; col5a1^−/−^; kdrl:EGFP).* Data are graphed as mean ± SEM. Statistics: Mann-Whitney *U* test. Asterisk representation: p-value < 0.05 (*); p-value < 0.005 (**); p-value < 0.0001 (****). Scale bars: (A) 250 μm; (B,E) 100 μm; (F) 50 μm.

To further characterize the hemorrhage phenotype in *collagen* mutants, we developed a phenotype screening procedure (Fig. 6C). In this assay, we analyzed both single mutants as well as double mutants to test genetic interactions between *col1a2* and *col5a1.* In addition, we tested whether the increase in physical stress exacerbates the hemorrhage phenotype by raising embryos in a high-viscosity medium (0.6% methylcellulose, MC) (56). Briefly, heterozygous adults harboring either one or both of the *collagen* mutations *(col1a2^+/−^, col5a1^+/−^,* or *col1a2^+/^-; col5a1^+/−^*) were intercrossed to generate wild type, heterozygous, and homozygous sibling embryos for different combinations (Fig. 6C). The resulting embryos were then incubated in either water or the viscous methylcellulose solution from 48 to 80 hpf, during which embryos were imaged and screened for hemorrhage in the trunk. Embryos were then genotyped to correlate the genotype with the phenotype. Similar to their wild-type *col1a2^+/+^* siblings (water: 0/14; MC: 0/27) and heterozygous *col1a2^+/−^* siblings (water: 0/39; MC: 0/38), homozygous *col1a2^−/−^* mutants did not show any trunk hemorrhage in either water (0/22) or the methylcellulose solution (0/21). This result suggests that loss of *col1a2* alone does not compromise vascular stability even with increased physical stress. By contrast, while neither wild-type *col5a1^+/+^* embryos (0/11) nor heterozygous *col5a1^+/−^* siblings (0/39) showed any hemorrhage under normal conditions, 12% of *col5a1^−/−^* embryos (3/25) displayed obvious hemorrhages in the trunk at 2 dpf, which was worsened substantially by physical stress (17/26, 65%). Analysis of different allele combinations of double mutants revealed several key characteristics of the hemorrhage phenotype (Fig. 6D). First, the loss of both *col5a1* wild-type alleles was required to cause the trunk hemorrhage phenotype. Second, even though loss of *col1a2* alone in *col1a2^+/−^* or *col1a2-^/-^* embryos was not sufficient to cause hemorrhage, loss of one or two *col1a2* wild-type alleles progressively increased the penetrance of the hemorrhage phenotype in *col5a1^−/−^* mutants, from 23% in *col1a2^+/−^; col5a1^−/−^* embryos (5/22) to 58% in *col1a2-^/-^; col5a1^−/−^* embryos (7/12) (Fig. 6D). Lastly, increased physical stress by incubating embryos in the viscous methylcellulose solution substantially exacerbated both the penetrance and the severity of the hemorrhage phenotype (Fig. 6D, 6E and S6D). It is interesting to note that stress-induced blood vessel rupture is typical of the vascular phenotypes associated with classical Ehlers-Danlos syndrome in humans. Together, our mutant analysis suggests that Col5a1 and Col1a2 function redundantly to maintain blood vessel integrity with Col5a1 being a more critical component.

The distribution of hemorrhagic foci is consistent with defects in ISVs in the trunk. To directly examine the ISV morphology in *collagen* mutants, we crossed *col5a1^+/−^; kdrl:EGFP* adults with *col1a2^+/−^; col5a1^+/−^* double heterozygous fish. Resulting embryos were incubated in the methylcellulose solution and their ISVs were analyzed at 53 hpf. In contrast to wild-type siblings, some *col5a1^−/−^* mutants showed severe ISV deformity and breakage, accompanied by hemorrhage (Fig. 6F). Quantifications of the diameter variability of intact ISVs as described earlier showed that both *col1a2^+/+^; col5a1^−/−^* and *col1a2^+/−^; col5a1^−/−^* embryos displayed significantly more variable ISVs than wild-type siblings (Fig. 6G). Consistent with a stronger hemorrhage phenotype, ISVs in *col1a2^+/−^; col5a1^−/−^* embryos were significantly more variable than those in *col1a2^+/+^*; *col5a1^−/−^* single mutants. Together, our work suggests that the deposition of Col5a1 and Col1a2 between 1-2 dpf, likely by perivascular fibroblasts, is crucial for the stabilization of nascent ISVs.

## DISCUSSION

In this study, we characterize the origin and function of perivascular fibroblasts, a novel population of blood vessel associated cells in zebrafish. Our work provides three main conclusions. First, perivascular fibroblasts originate from the sclerotome and are distinct from pericytes. Second, a subset of perivascular fibroblasts functions as pericyte progenitors. Third, cell ablation and mutant analysis reveal that perivascular fibroblasts stabilize nascent blood vessels through Col1a2 and Col5a1. Together, we propose a dual role of perivascular fibroblasts in vascular stabilization: they first establish the ECM around nascent blood vessels for initial stabilization and then function as pericyte progenitors to generate pericytes that further contribute to vessel integrity (Fig. 7). Our work provides new insights into the molecular and cellular mechanisms underlying vascular stabilization and vascular diseases, such as Ehlers-Danlos syndrome.

**Fig. 7.**
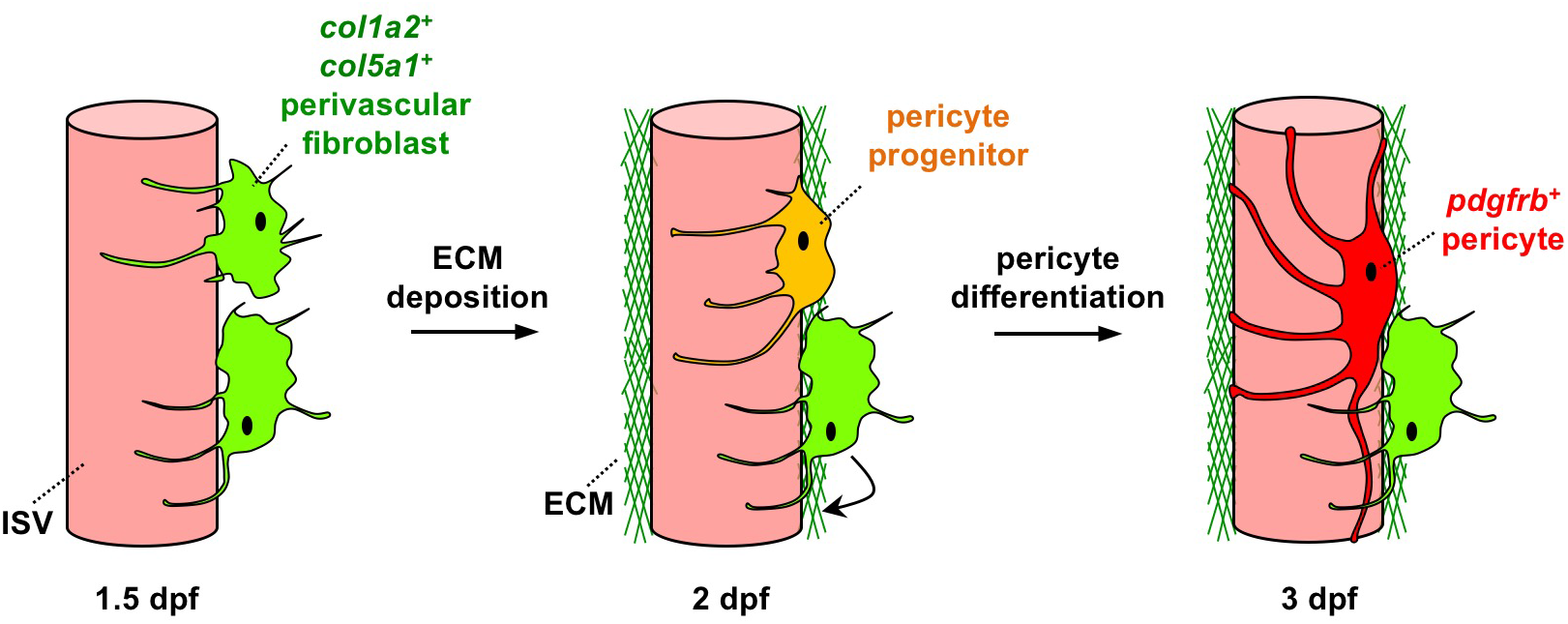
Model of perivascular fibroblasts in vascular stabilization in zebrafish. Perivascular fibroblasts, characterized by the expression of fibrillar collagens *col1a2* and *col5a1,* become associated with intersegmental vessels (ISVs) in the zebrafish trunk by 1.5 dpf. Perivascular fibroblasts deposit a network of collagen fibers to establish the vascular ECM and stabilize nascent ISVs at 2 dpf and continue to deposit collagen until at least 3 dpf. In the same time window, a subset of perivascular fibroblasts functions as pericyte progenitors. They gradually upregulate the expression of the classic pericyte marker *pdgfrb*, and develop elongated cellular processes. By 3 dpf, these ‘pericyte progenitors’ have completely differentiated into mature pericytes, showing robust *pdgfrb* expression distinct from *col1a2^+^* perivascular fibroblasts. Together, perivascular fibroblasts perform dual functions in vascular stabilization by depositing collagens to support nascent blood vessels and acting as pericyte progenitors.

### Perivascular fibroblasts are a unique population of perivascular cells

Mural cells are well studied blood vessel support cells. In the zebrafish trunk, each intersegmental vessel is associated with 1-2 pericytes at 4 dpf. Four lines of evidence suggest that collagen-expressing perivascular fibroblasts represent a unique population of perivascular cells, distinct from pericytes. First, these two cell populations show distinct marker expression, with pericytes labeled by *pdgfrb,* and perivascular fibroblasts marked by the expression of collagen genes, such as *col1a2* and *col5a1.* Second, perivascular fibroblasts become associated with ISVs as early as 31 hpf, whereas pericytes do not differentiate until at least 60 hpf (45). Third, perivascular fibroblasts are much more abundant than pericytes, with on average 10.6 perivascular fibroblasts per ISV compared to 0.9 pericytes per vessel at 4 dpf. Finally, perivascular fibroblasts and pericytes display vastly distinct morphologies. Similar to mammalian pericytes, ISV pericytes in zebrafish have small and flat cell bodies with long cellular processes that tightly wrap around and along the underlying endothelium. By contrast, perivascular fibroblasts are more loosely associated with ISVs, and appear more globular with numerous shorter and thinner processes that extend around the endothelium. Based on the differences in marker expression, developmental timing, cell number, and cell morphology, we propose that perivascular fibroblasts represent a novel perivascular cell population, distinct from pericytes. Intriguingly, recent single cell RNA sequencing studies have revealed a similar perivascular ‘fibroblast-like’ cell population in the adult murine brain (Saunders et al., 2018; Vanlandewijck et al., 2018). Similar to zebrafish perivascular fibroblasts, murine fibroblast-like cells are loosely adhered to the blood vessels, abluminal to mural cells and the basement membrane. Furthermore, murine fibroblast-like cells robustly express many ECM genes, including collagen genes *col5a1* and *col1a2,* but not mural cell markers such as *pdgfrb.* Combined with our work, these studies suggest that perivascular fibroblasts represent a unique and evolutionarily conserved population of blood vessel associated cells across vertebrates.

### The sclerotome is the embryonic origin of perivascular fibroblasts

The sclerotome is the embryonic compartment that contributes to the axial skeleton of vertebrates. Our previous and current studies demonstrate that the sclerotome also gives rise to several populations of tissue support cells, including tenocytes (tendon fibroblasts) (46) and perivascular fibroblasts. The zebrafish sclerotome has a unique bipartite organization with a larger ventral domain and a smaller dorsal domain. Using time-lapse imaging and cell tracing, we have built a spatial map of origin for perivascular fibroblasts. Similar to tenocytes, the ventral sclerotome domain is the main contributor of perivascular fibroblasts (84%). Furthermore, different sclerotome domains contribute to perivascular fibroblasts with a stereotypic spatial pattern. The ventral sclerotome domain generates perivascular fibroblasts along the entire length of the ISV, whereas the dorsal sclerotome domain contributes only locally to perivascular fibroblasts associated with the dorsal half of the ISV. Since perivascular fibroblasts also function as pericyte precursors (see below), our results suggest that most pericytes associated with ISVs also originate from the sclerotome. Consistent with our results, vSMCs around the dorsal aorta have been traced back to cells derived from the sclerotome in zebrafish (50). In addition, mural cells in the zebrafish trunk vessels have been shown to be dependent on the mesoderm but not the neural crest lineage (57). Thus, blood vessel support cells in the zebrafish trunk, including perivascular fibroblasts and mural cells, likely originate from the sclerotome. Combined with previous lineage analysis in mouse and chick (58, 59), these data suggest a model where mural cells in the brain are neural crest derived, whereas mural cells in the trunk are sclerotome derived.

### Perivascular fibroblasts function as pericyte progenitors

Although perivascular fibroblast-like cells have been identified in the mouse brain by single cell RNA sequencing, their biological functions are still unclear. Two complementary experiments suggest that perivascular fibroblasts function as pericyte progenitors. First, in vivo imaging reveals that the majority of newly born pericytes on ISVs at 3 dpf can be traced back to perivascular fibroblasts at 2 dpf. Indeed, time-lapse imaging in both *nkx3.1^NTR-mCherry^; pdgfrb:GFP* and *col1a2:GFP; pdgfrb^NTR-mCherry^* embryos reveals similar kinetics of pericyte differentiation: a few perivascular fibroblasts gradually downregulate their fibroblast marker expression, upregulate pericyte reporters, and undergo morphological changes to extend long pericyte-like processes along ISVs at 3 dpf. In complementary experiments, genetic ablation of perivascular fibroblasts results in a 3.5-fold reduction in the number of pericytes. Together, these results indicate that some perivascular fibroblasts serve as pericyte precursors, contributing to most, if not all, pericytes associated with ISVs in the trunk. Incomplete labeling or ablation likely reflects the mosaic nature of the *col1a2* transgenic lines (49).

It is interesting to note that there are about 10 perivascular fibroblasts associated with each ISV, however, only 1-2 of these cells differentiate into pericytes. It thus raises the question of how these perivascular fibroblasts are ‘selected’ as pericyte progenitors. There are two possibilities. In the first scenario, all perivascular fibroblasts have the equal potential to differentiate into pericytes, but extrinsic cues determine which cell to switch on the pericyte fate. Notch signaling has been implicated in the regulation of pericyte formation in both mouse and zebrafish (57; 60–63). In particular, recent work in zebrafish has shown that the activation of Notch signaling in naïve mesenchymal cells (likely corresponding to perivascular fibroblasts in our study) is crucial to induce pericyte differentiation around ISVs (Ando et al., 2019). Therefore, active Notch signaling might bias some perivascular fibroblasts towards the pericyte lineage. Alternatively, perivascular fibroblasts might represent a heterogenous population of cells, containing a small fraction of pericyte progenitors. Supporting this idea, recent scRNA sequencing studies have shown that perivascular fibroblast-like cells in the mouse brain can be further divided into several subtypes based on unique gene expression signatures (Saunders et al., 2018; Vanlandewijck et al., 2018).

### Perivascular fibroblasts function to stabilize nascent blood vessels

Although mural cells are known to stabilize mature blood vessels, it is not clear how newly formed blood vessels are supported prior to the differentiation of mural cells. Our work provides strong evidence that perivascular fibroblasts function to stabilize nascent blood vessels by secreting collagens. First, time-lapse imaging reveals that the emergence of perivascular fibroblasts along ISVs occurs concurrently with ISV development, at least one day before the differentiation of the first pericytes. Second, genetic ablation of perivascular fibroblasts prior to pericyte differentiation results in aberrant ISV morphology. By contrast, late ablation of perivascular fibroblasts from 4 to 5 dpf has no effects on the ISV morphology. These results suggest that perivascular fibroblasts play a crucial role in supporting newly formed blood vessels before pericyte differentiation. Lastly, perivascular fibroblasts express high level of collagen genes, such as *col1a2* and *col5a1,* and contribute to the vascular collagen surrounding ISVs. The live Col1a2-GFP reporter reveals that perivascular fibroblasts contribute to a network of collagen fibers around nascent ISVs as early as 2 dpf, which continues to expand until at least 3 dpf. Strikingly, loss of *col5a1* results in dysmorphic ISVs with spontaneous hemorrhage in the trunk, which is exacerbated by the additional loss of *col1a2* or increasing physical stress. It is important to note that these ISV phenotypes can be observed as early as 2 dpf, well before the emergence of pericytes on ISVs. Together, our experiments show that perivascular fibroblasts act to stabilize nascent ISVs prior to pericyte differentiation, likely through regulating collagen deposition. Interestingly, ablation of perivascular fibroblasts rarely results in the severe hemorrhage phenotype observed in *col5a1^−/−^* mutants (under either normal or stressed conditions). One possible explanation is that the few perivascular fibroblasts spared from the ablation due to transgene mosaicism might be sufficient to prevent ISV rupture. Consistent with our work, recent study has described a similar population of *col22a1^+^* perivascular fibroblast-like cells associated with the cranial vasculature in zebrafish (64). Null mutations in *col22a1,* a minor fibril-associated collagen, result in compromised cranial vessel integrity in both embryos and adults following cardiovascular stress (64), highlighting the importance of collagen and perivascular fibroblasts in vascular stabilization.

### Zebrafish model of Ehlers-Danlos syndrome

Collagen is the most abundant protein in the human body and is a major component of the ECM (65). Fibrillar collagens, including collagen I and collagen V, constitute the most common sub-group within the collagen family which often co-localize in tissues such as the dermis, tendons and ligaments (65). Mutations in fibrillar collagens or collagen modification enzymes have been implicated in a number of connective tissue diseases, including Ehlers-Danlos syndrome (EDS). The classical subtype of EDS, caused by mutations in collagen V, and less frequently collagen I, is characterized by skin hyperextensibility, joint hypermobility, and blood vessel fragility (19,66). Classical EDS patients experience easy bruising and excessive hemorrhaging under normal exertion. By contrast, vascular phenotypes are more prominent in the vascular subtype of EDS, resulting from mutations in collagen III, with most patients experiencing severe spontaneous rupture of large arteries (19). Interestingly, the zebrafish genome does not contain a collagen III gene, which is likely lost during evolution as in many other teleost fish species (67). Characterization of zebrafish *col1a2* and *col5a1* mutants reveals several key features of vascular EDS. Embryos lacking functional *col5a1* genes develop spontaneous hemorrhages in the trunk under normal physiological conditions, reminiscent of easy bruising in patients with vascular EDS. One hallmark of vascular EDS is that physical stress often leads to more severe symptoms (68). Similarly, both the penetrance and severity of the hemorrhage phenotype in *col5a1^−/−^* mutants are strongly enhanced by the increase of physical stress. Thus, our results suggest that the lack of collagen III in zebrafish is likely compensated by other fibrillar collagens (types I and V). Our *col1a2* and *col5a1* mutants represent the first zebrafish model of vascular EDS that we know of.

Analysis of collagen mutants also provides new insights on vascular EDS. First, Col5a1 is a more critical component of the vascular ECM compared to Col1a2. Lack of *col5a1* function leads to ISV hemorrhages, whereas loss of *col1a2* alone does not result in any obvious vascular defects even with increased physical stress. Intriguingly, collagen I is the most abundant component in collagen fibrils (> 90%), while collagen V constitutes only a minor fraction (< 5%) (69). Why do *col5a1^−/−^* mutants show more severe phenotypes than *col1a2^−/−^* mutants? Previous work has shown that Col5a1 plays an important role in proper nucleation and diameter regulation of collagen fibrils (70). The loss of Col5a1 might result in defects in the assembly of collagen I into fibrils, leading to a more severe phenotype. Second, our mutant analysis reveals genetic interactions between *col1a2*and *col5a1.* Loss of wildtype *col1a2* alleles in the *col5a1^−/−^* background substantially enhances both the penetrance and the severity of the hemorrhage phenotype in a dosage dependent manner. This result suggests that Col5a1 and Col1a2 function redundantly to maintain vascular integrity. Lastly, as discussed above, our work provides strong evidence that defects in perivascular fibroblasts are the underlying cellular basis for the vascular phenotypes of EDS patients. Perivascular fibroblasts thus represent the cellular targets for potential therapeutic interventions.

In summary, our work identifies perivascular fibroblasts as a novel perivascular cell population with dual function in vascular stabilization: they regulate the ECM of nascent blood vessels and also function as pericyte progenitors. Interestingly, many perivascular fibroblasts remain associated with mature blood vessels even after pericyte differentiation. This raises several questions about the function of these cells at later stages. Do perivascular fibroblasts continue to regulate the ECM around the mature vasculature? Do they function as resident stem cells to replace aging/damaged pericytes? Previous work has suggested that tissue injury in the mouse brain and spinal cord often triggers a fibrotic response by blood vessel associated cells (25,34–36). Attenuation of the fibrotic response has been shown to limit scar formation and improve axon regeneration post spinal cord injury in mice, suggesting that perivascular stromal cells might be an important therapeutic target (71). It is plausible that zebrafish perivascular fibroblasts also play an active role in tissue injury repair, an exciting question for future investigations.

## MATERIALS AND METHODS

### Zebrafish strains

Zebrafish strains were maintained according to standard protocols. Animal research was conducted in accordance with current guidelines of the Canadian Council on Animal Care. All protocols were approved by the Animal Care Committee at the University of Calgary (#AC17-0128). The following transgenic strains were utilized in this study: *TgBAC(col1a2:Gal4)ca102* (46, 49), *TgBAC(col1a2:GFP)ca103* (46), *Tg(kdrl:EGFP)la116* (72), *Tg(kdrl:mCherry)ci5* (73), *TgBAC(nkx3.1:Gal4)ca101* (46), *TgBAC(pdgfrb:Gal4FF)ca42* (45,74), *TgBAC(pdgfrb:GFP)ca41* (45), *Tg(UAS:Kaede)s1999t* (75), *Tg(UAS:NTR-mCherry)c264* (75). The mosaic *col1a2:Gal4; UAS:Kaede* line was maintained by growing embryos with more mosaic Kaede expression. The *col1a2^ca108^* and *col5a1^ca109^* mutant lines were maintained as heterozygotes, and homozygous embryos were generated by intercrossing heterozygous carriers.

### Generation of CRISPR mutants

The *col1a2^ca108^* and *col5a1^ca109^* mutant lines were generated using the CRISPR/Cas9 system as previously described (53). Briefly, target sites were identified using the web program CHOPCHOP (http://chopchop.cbu.uib.no) (76). sgRNA target sequences are 5’-GGGGGTTCCATTTGATCCAG-3’ *(col1a2*) and 5’-GGCTCCAGCAGATCATCCAG-3’ *(col5a1).* To assemble DNA templates for sgRNA transcription, gene-specific oligonucleotides containing the T7 promoter sequence (5’-TAATACGACTCACTATA-3’), the 20 base target site, and a complementary sequence were annealed to a constant oligonucleotide encoding the reverse-complement of the tracrRNA tail. sgRNAs were generated by in vitro transcription using the Megascript kit (Ambion). Cas9 mRNA was transcribed from linearized pCS2-Cas9 plasmid using the mMachine SP6 kit (Ambion). To generate mutants, onecell stage wild-type embryos were injected with a mix containing the appropriate sgRNA at 20 ng/μl and Cas9 mRNA at 200 ng/μl. Injected fish were raised to adulthood and crossed to generate F1 embryos. T7 Endonuclease I assay (NEB) was then used to identify the presence of indel mutations in the targeted region of F1 fish. A *col1a2* allele containing 1 bp deletion *(col1a2^ca108^*) and a *col5a1* allele with 4 bp deletion *(col5a1^ca109^*) were identified (Fig. S5). For genotyping, small regions around the mutation sites were specifically amplified by PCR, which was followed by allele-specific restriction enzyme analysis. For *col1a2^ca108^,* the primers used were 5’-TTTTAAAGACTCACATTTGCCTT-3’ (forward) and 5’-CTCCGGGCTAGCTTTATATTTCGATT-3’ (reverse). For *col5a1^ca109^,* the primers used were 5’-ACTCTTGTTTGCTGTGCAGGT-3’ (forward) and 5’-CTCACCGGTATTGGCCGTGTT-3’ (reverse). Following PCR, the products were digested with a restriction enzyme which cuts only the wild type or the mutant allele. The enzymes used were BclI and BstCI (NEB) for *col1a2^ca108^* and *col5a1^ca109^,* respectively. The resulting band sizes were analyzed using gel electrophoresis. For *col1a2^ca108^,* the BclI digest resulted in a single band of 473 bp in wild-type embryos, and two bands of 400 bp and 73 bp in homozygous *col1a2^−/−^* embryos. For *col5a1^ca109^,* the BstCI digest resulted in multiple bands at 89, 44, 30, 18 and 9 bp in wild-type embryos, and bands of 107, 44, 30 and 9 bp in homozygous *col5a1^−/−^* embryos.

### Plasmid injection

To visualize collagen distribution, we generated a *UAS:Col1a2-GFP* construct based on a previously published *pME-Col1a2-GFP* plasmid (52). *UAS:Col1a2-GFP* (40 ng/μl) was co-injected with *tol2* transposase mRNA (40 ng/μl) into *nkx3.1:Gal4; UAS:NTR-mCherry* embryos at the one-cell stage with 1 nl per embryo. For more mosaic labeling, *UAS:Col1a2-GFP* was injected at 10 ng/μl. Injected embryos were screened for GFP expression at appropriate stages for imaging.

### In situ hybridization and immunohistochemistry

Whole-mount in situ hybridization and antibody staining were performed according to standard protocols. *col1a2* and *col5a1* antisense probes were used in this study. Double fluorescent in situ hybridization was performed using digoxigenin (DIG) and dinitrophenyl (DNP) labeled probes. For antibody labeling, the rabbit polyclonal antibody to GFP (1:500, MBL) was used. For fluorescent detection of antibody labeling, appropriate Alexa Fluor-conjugated secondary antibodies were used (1:500, Thermo Fisher).

### Time-lapse imaging

Imaging was performed using the Olympus FV1200 confocal microscope as previously described (46). Embryos older than 24 hpf were incubated in fish water with 1-phenyl 2-thiourea (PTU) to prevent pigmentation. Fish were anesthetized in 0.4% tricaine and mounted in 0.6% low melting point agarose. Z-stack images of the region of interest were then collected using the 20x objective at appropriate time intervals (5-15 mins) for up to 24 hours. Movies and images were processed using the Olympus Fluoview software and the Fiji software (77). Cell tracing was performed manually using Fiji.

### Cell ablation

To ablate perivascular fibroblasts, *nkx3.1^NTR-mCherry^* or *col1a2^NTR-mCherry^* embryos at desired developmental stages were treated with water (control group) or 5 mM metronidazole (MTZ, experimental group) for 24-48 hours. In some experiments, mCherry^−^ siblings treated with 5 mM MTZ in the same time windows were used as additional controls. Embryos were subsequently washed in fish water and imaged to confirm successful ablation of mCherry^+^ cells.

### Quantification of blood vessel diameter variability

To quantify blood vessel diameters, we used the endothelial specific *kdrl:GFP* reporter to fluorescently label intersegmental vessels. Embryos were imaged laterally in the trunk region (somite 12 to 18) and 8-10 ISVs were quantified per embryo. Using the ‘line selection’ tool in Fiji, four diameter measurements were taken for each ISV at equidistant points along its length and measurements were averaged to obtain mean vessel diameter. Standard deviation from the mean vessel diameter was plotted as a readout of ISV diameter variability.

### Mechanical stress assay

To test the effect of physical stress on the vascular phenotype of collagen mutants, we adopted a mechanical overloading assay by raising embryos in a high-viscosity medium (0.6% methylcellulose) (56). Heterozygotes for single mutants *(col1a2^+/−^*or *col5a1^+/−^*) or double mutants *(col1a2^+/−^; col5a1^+/−^*) were intercrossed to obtain sibling embryos with different allele combinations. Resulting embryos were incubated in water (control group) or 0.6% methylcellulose (MC, experimental group) at 48 hpf. Fish were screened for hemorrhage in the trunk region approximately every 3 hours from 48 to 80 hpf and subsequently genotyped for *col1a2* and *col5a1.*

### Statistical analysis

All graphs and statistical analysis were generated using the GraphPad PRISM software. Data were plotted with mean ± SEM indicated. Significance was calculated by performing using the nonparametric Mann-Whitney *U* test with two-tailed p values: p > 0.05 (ns, not significant), p < 0.05 (*), p < 0.01 (**), p < 0.001 (***) and p < 0.0001 (****).

## ACKNOWLEDGEMENTS

We thank the zebrafish community for providing probes and reagents; Paul Martin for sharing the *Col1a2-GFP* plasmid; Sarah Childs for sharing transgenic lines and providing critical input on this project; members of the Childs and Huang laboratories for discussions; and Paul Mains for critical comments on the manuscript.

## SUPPLEMENTAL FIGURES

**Fig. S1.**
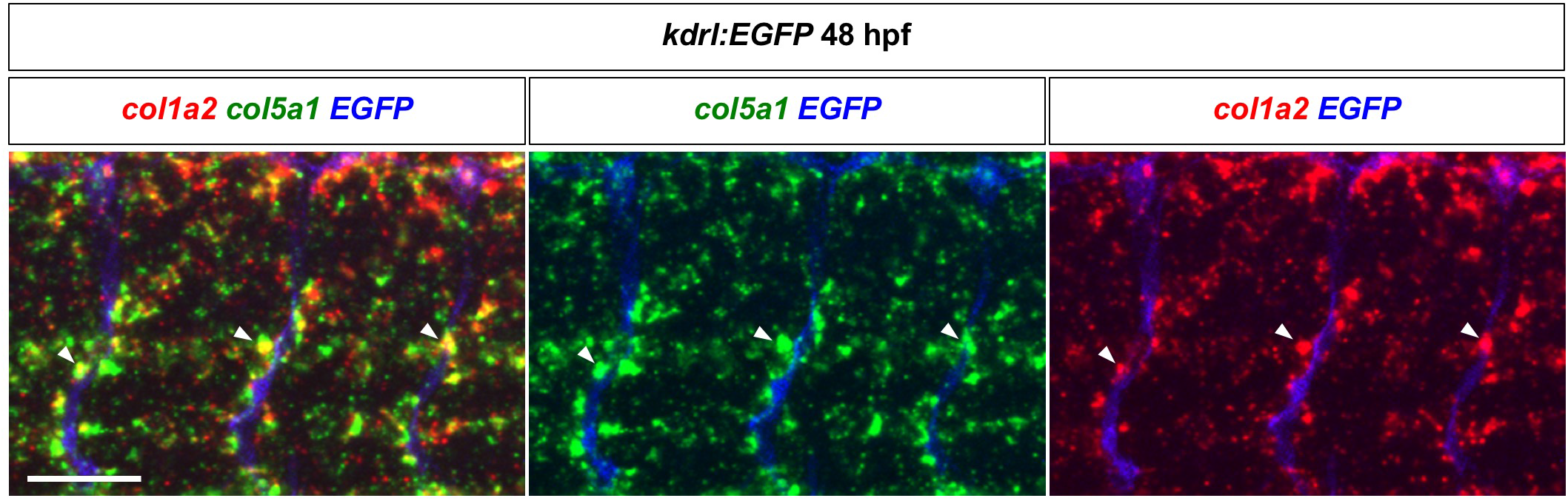
Co-expression of *col1a2* and *col5a1* in perivascular fibroblasts. *kdrl:EGFP* embryos at 48 hpf were co-labeled with *col1a2* (red) and *col5a1* (green) by double fluorescent in situ hybridization followed by immunofluorescence labeling using the GFP antibody (blue). Co-expression of *col1a2* and *col5a1* is observed in perivascular fibroblasts (arrowheads) along EGFP^+^ ISVs. *n* = 23 embryos. Scale bar: 50 μm.

**Fig. S2.**
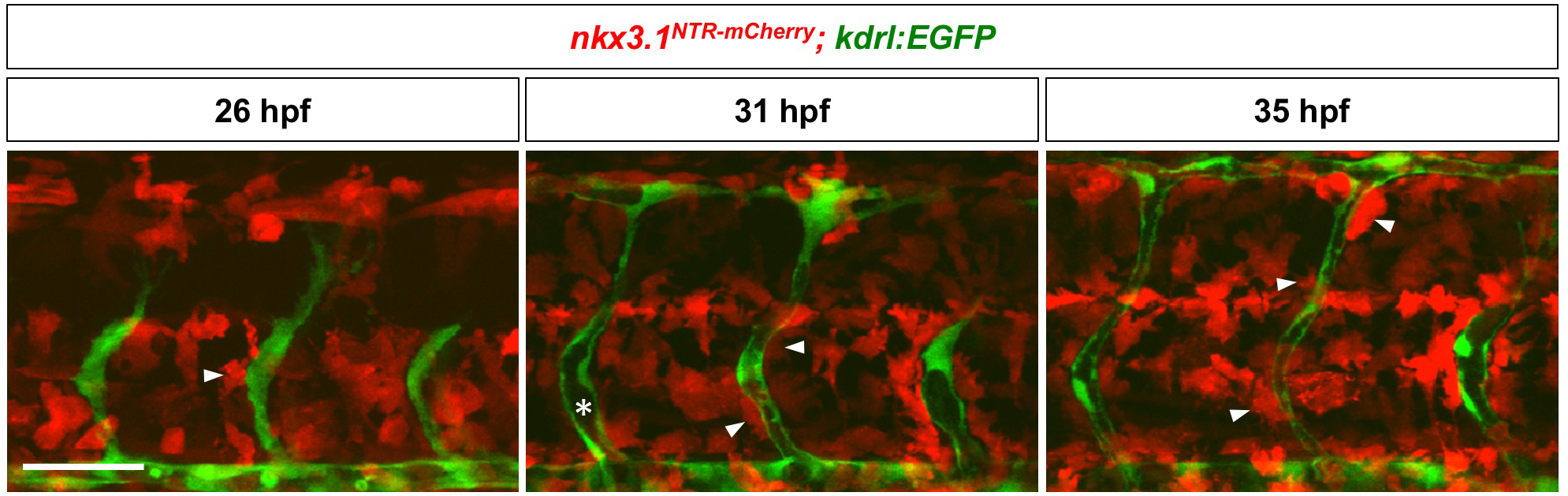
Developmental time-course of perivascular fibroblasts. *nkx3.1^NTR-mCherry^; kdrl:EGFP* embryos were imaged at 26, 31, and 35 hpf to visualize different stages of perivascular fibroblast development. At 26 hpf, some *nkx3.1^NTR-mCherry^* cells (arrowhead) were visible along the ventral half of ISV sprouts. As ISV lumenization became visible (asterisk) at 31 hpf, more *nkx3.1^NTR-mCherry^* cells (arrowheads) appeared along ISVs. By 35 hpf, mCherry^+^ cells (arrowheads) were present along the entire length of ISVs. Scale bar: 50 μm.

**Fig. S3.**
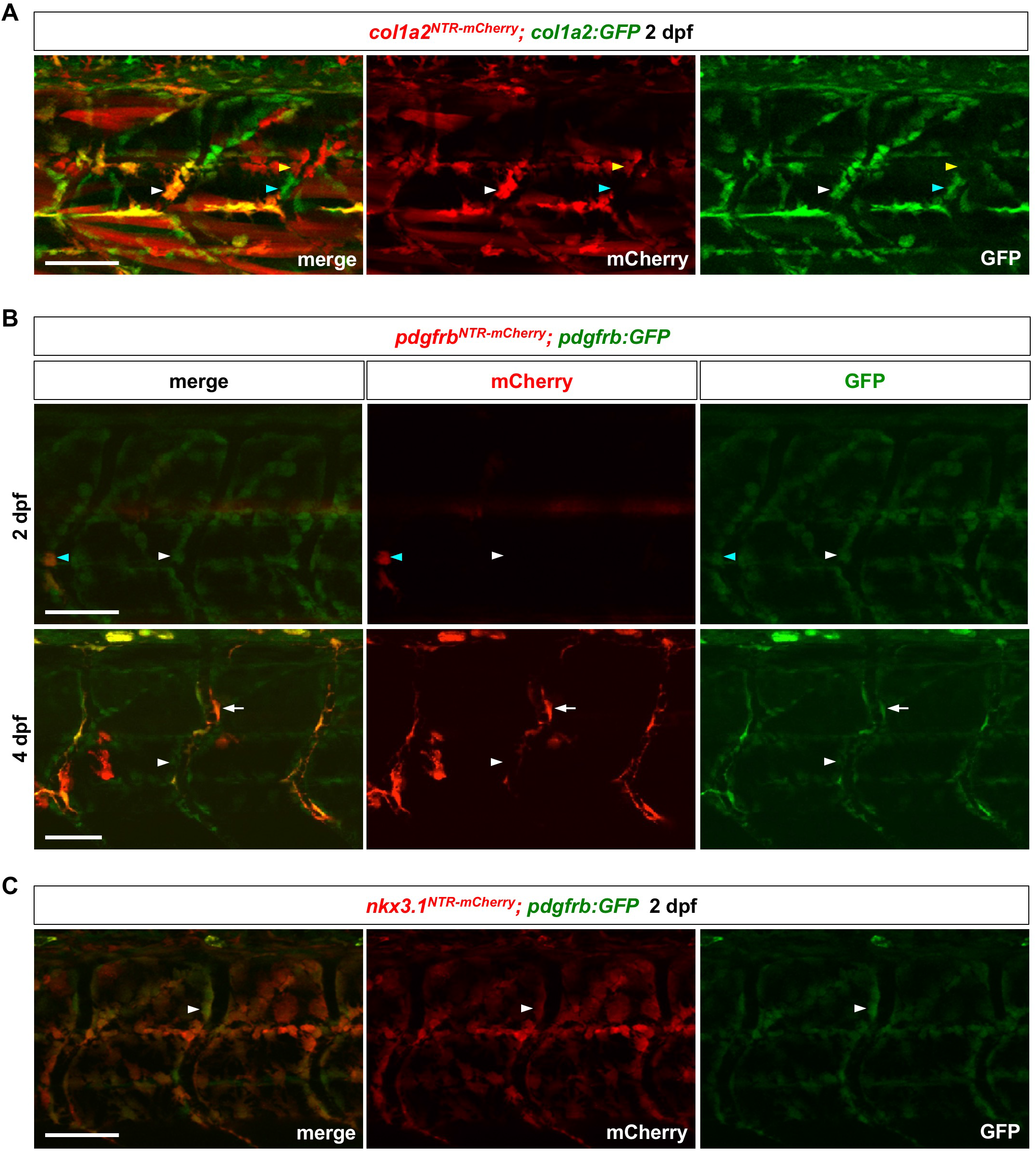
Characterization of transgenic reporters. (A) *col1a2^NTR-mCherry^; col1a2:GFP* embryos were imaged at 2 dpf. Due to the mosaic nature of both reporters, some perivascular fibroblasts were GFP^+^mCherry^+^ (white arrowheads), some GFP^+^mCherry^−^ (cyan arrowheads), and some GFP^−^ mCherry^+^ (yellow arrowheads). *n* = 5 embryos. (B) *pdgfrb^NTR-mCherry^; pdgfrb:GFP* embryos were imaged at 2 dpf (top panel) and 4 dpf (bottom panel). At 2 dpf, most perivascular fibroblasts were GFP^+^mCherry^−^ (white arrowheads) while a few cells were GFP^+^mCherry^+^ (cyan arrowheads). At 4 dpf, pericytes were GFP^high^mCherry^+^ (arrows) whereas perivascular fibroblasts were GFP^low^mCherry^−^ (arrowheads). *n* = 15 embryos. (C) *nkx3.1^NTR-mCherry^; pdgfrb:GFP* embryos were imaged at 2 dpf. Perivascular fibroblasts (arrowheads) were positive for both *nkx3.1^NTR-mCherry^* (red) and *pdgfrb:GFP* (green) reporters. *n* = 15 embryos. Scale bars: 50 μm.

**Fig. S4.**
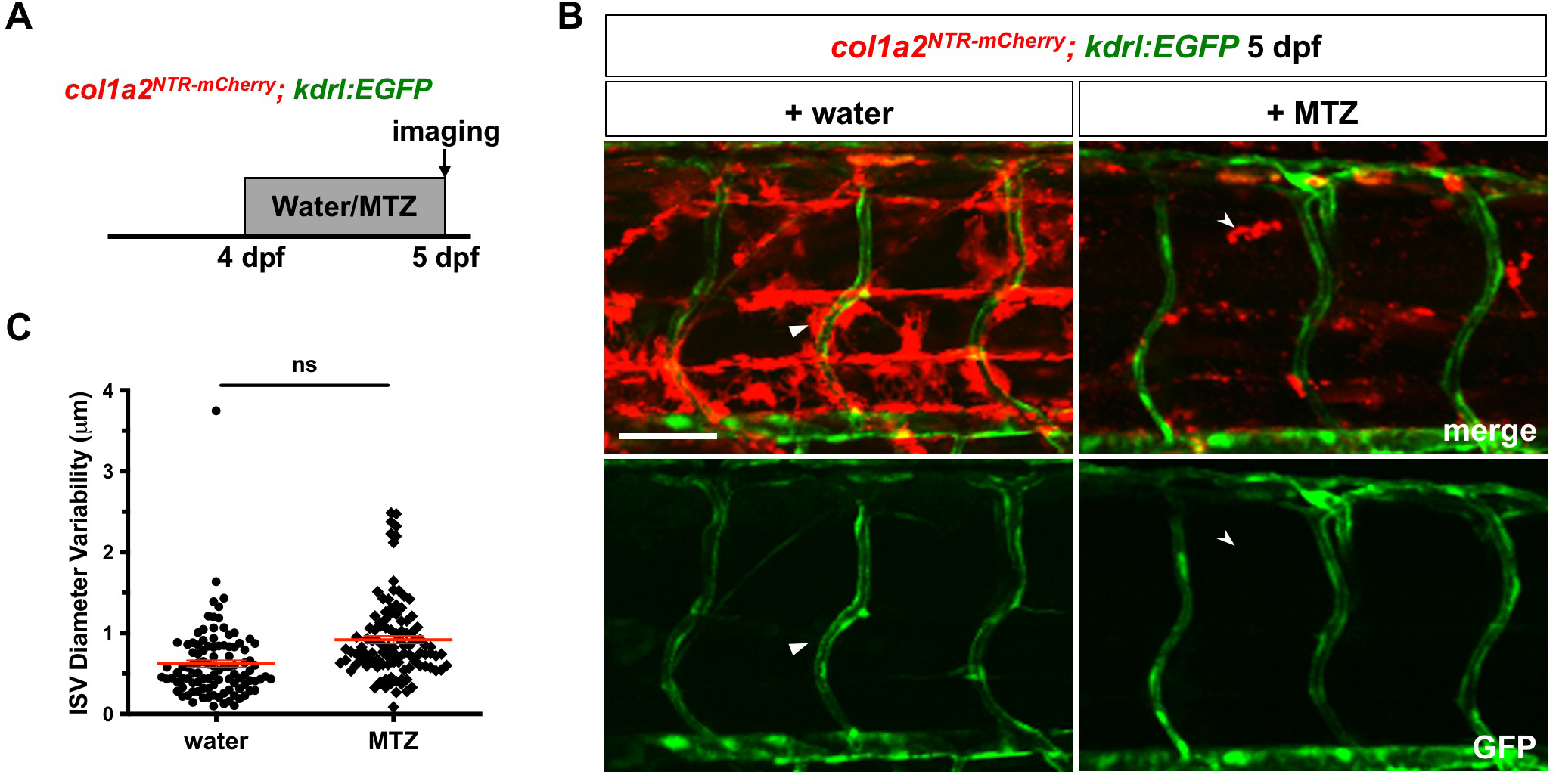
Late ablation of perivascular fibroblasts does not alter ISV morphology. (A) Schematic of experimental procedure for late perivascular fibroblast ablation. *col1a2^NTR-mCherry^; kdrl:EGFP* embryos were incubated in either water or MTZ from 4 to 5 dpf and imaged to visualize ISV morphology. (B) Representative images showing water (left) and MTZ (right) treated embryos. Water-treated control embryos had many mCherry^+^ cells (arrowheads), whereas MTZ treatment resulted in complete perivascular fibroblast ablation, with only mCherry^+^ debris visible (notched arrowheads). No distinguishable difference in ISV morphology was visible between MTZ treated and control embryos. (C) Quantification of ISV diameter variability in (B). ISV diameter and variability measurements were quantified as described in Fig 5. *n* = 103 ISVs from 9 embryos (water); 107 ISVs from 13 embryos (MTZ). Data are plotted as mean ± SEM. Statistics: Mann-Whitney *U* test. ns: not significant. Scale bar: (B) 50 μm.

**Fig. S5.**
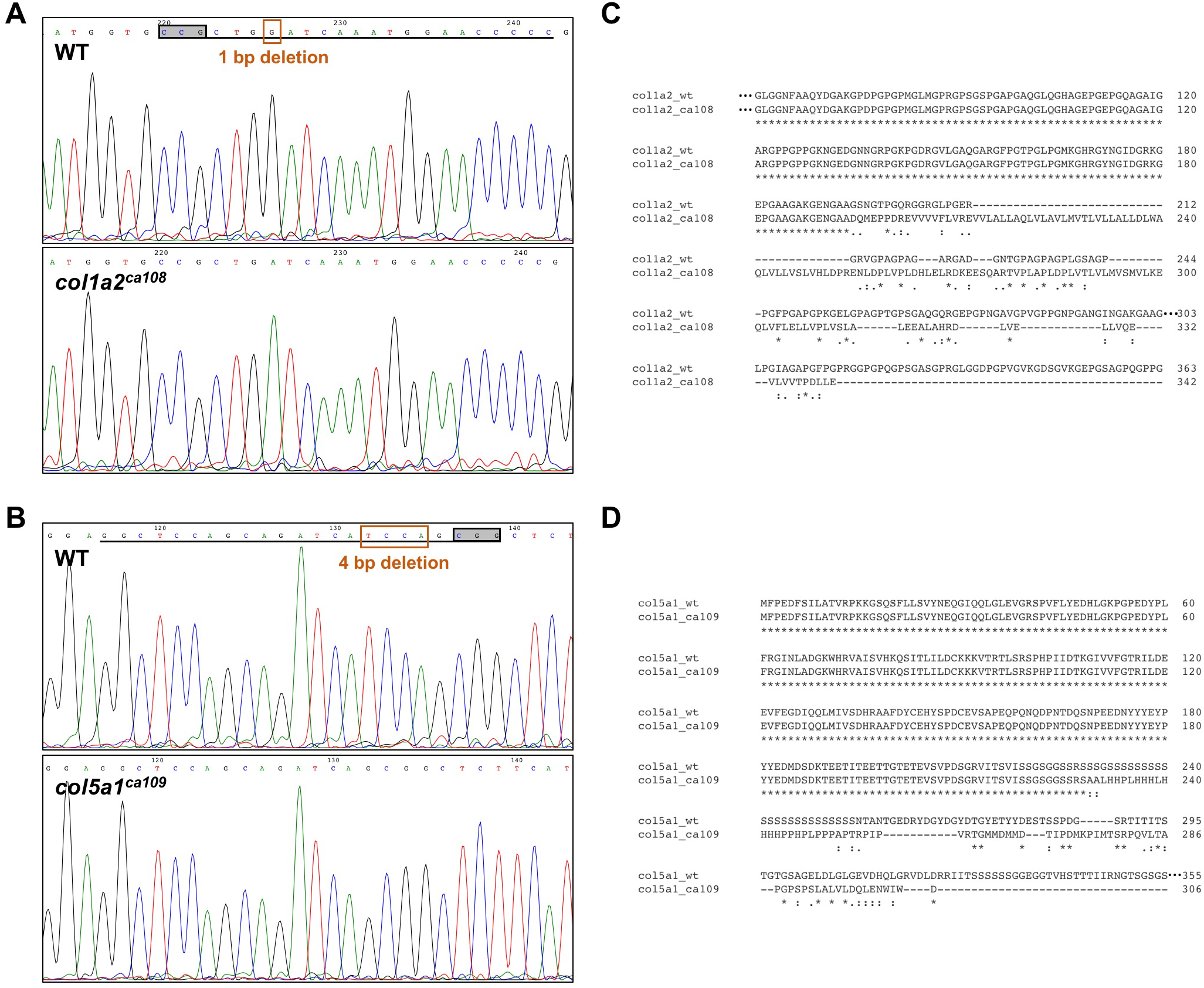
Generation of collagen mutants. (A) Sequencing chromatograms of *col1a2* WT and *col1a2^ca108^* sequences. The sgRNA target sequence is underlined and the PAM motif is highlighted with a black box. The 1bp deletion in the *col1a2^ca108^* sequence is denoted with an orange box. (B) Sequencing chromatograms of *col5a1* WT and *col5a1^ca109^* sequences. The sgRNA target sequence is underlined and the PAM motif is highlighted with a black box. The 4bp deletion in the *col5a1^ca109^* sequence is denoted with an orange box. (C) Alignment of *col1a2* WT and *col1a2^ca108^* protein sequences. (D) Alignment of *col5a1* WT and *col5a1^ca109^* protein sequences. Protein sequences were aligned using Clustal Omega (https://www.ebi.ac.uk/Tools/msa/clustalo/).

**Fig. S6.**
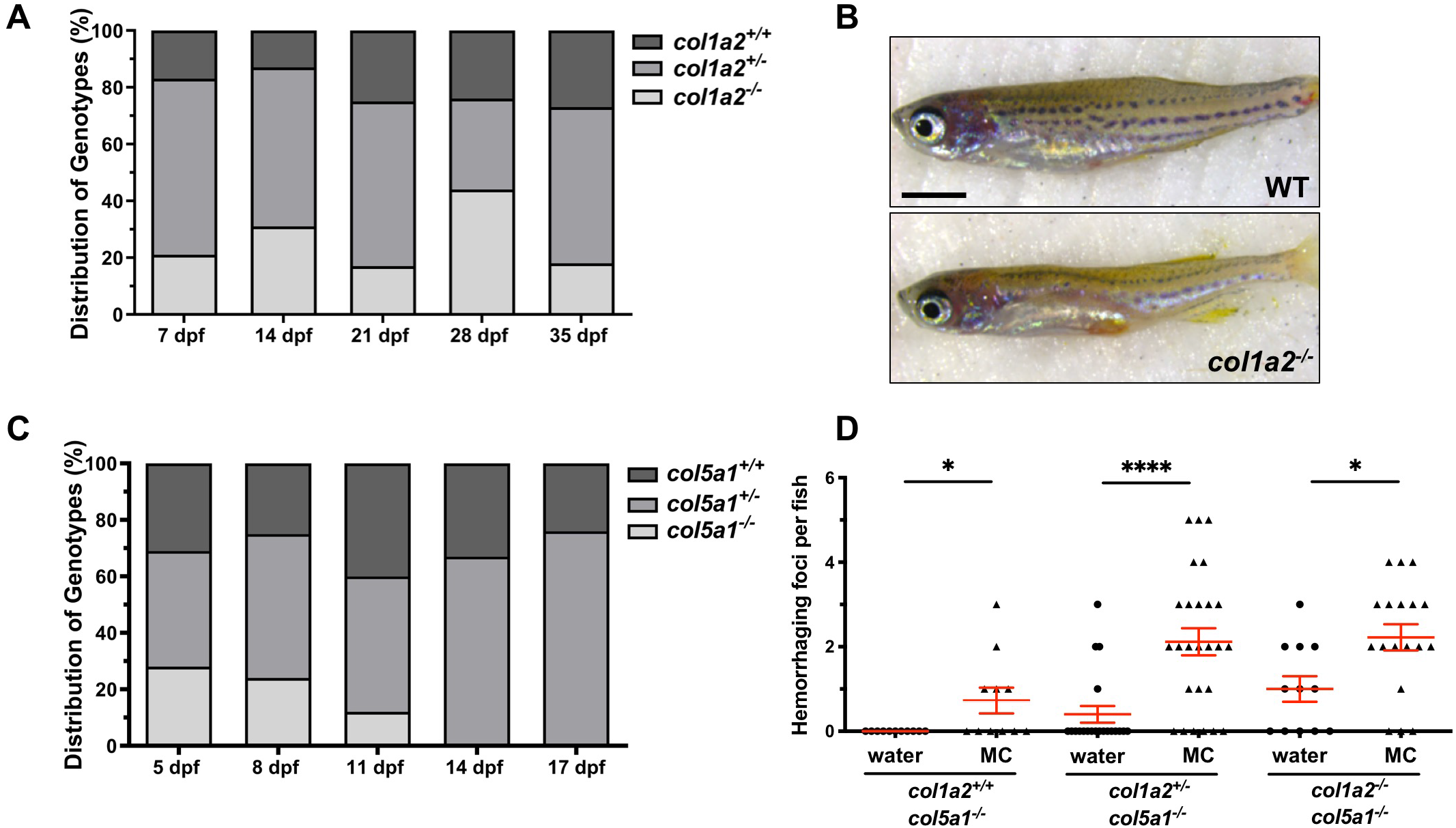
Characterization of collagen mutants. (A) Distribution of genotypes in progeny of *col1a2^+/−^* intercrosses. Embryos from the mentioned crosses were grown and genotyped every 7 days from 7 dpf to 35 dpf to determine genotype distribution. Distribution of genotypes followed roughly Mendelian ratios at all stages examined. (B) Comparison of adult wild type and *col1a2^−/−^* siblings. (C) Distribution of genotypes in progeny of *col5a1^+/−^* intercrosses. Embryos from crosses of *col5a1^+/−^* adults were grown and genotyped at 3 day intervals from 5 dpf to 17 dpf. While the distribution of genotypes followed mendelian ratios at 5 and 8 dpf, *col5a1^−/−^* fish were completely absent at 14 and 17 dpf. (D) Quantification of hemorrhage severity in collagen mutants shown in Fig 5D and 5E. Hemorrhage severity was scored by counting the number of visible hemorrhage foci present in the trunk. Fish with no visible hemorrhage were counted as 0. Increased physical stress in the viscous MC solution resulted in an increase in hemorrhage severity across mutants with different genotypes. *n* = 12 *(col1a2^+/+^; col5a1^−/−^* + water); 11 *(col1a2^+/+^; col5a1^−/−^* + MC); 20 *(col1a2^+/−^; col5a1^−/−^* + water); 26 *(col1a2^+/−^; col5a1^−/−^* + MC); 12 *(col1a2-^/-^; col5a1^−/−^* + water); and 18 *(col1a2^−/−^; col5a1^−/−^* + MC) embryos. Results were graphed as mean ± SEM. Statistics: Mann-Whitney *U* test. Asterisk representation: p-value < 0.05 (*); p-value < 0.0001 (****). Scale bar: (B) 2 mm.

**Movie S1. Perivascular fibroblasts originate from the sclerotome.** *nkx3.1^NTR-mCherry^; kdrl:EGFP* embryos were imaged from 25 hpf to 49.5 hpf at ~8 minute intervals (7 min 58 sec) with time stamps indicated in the hh:mm format. Perivascular fibroblasts along ISVs were retrospectively traced to determine their cell of origin. One cell from the ventral sclerotome domain (cyan arrows) and one cell from sclerotome derived notochord associated cells (yellow arrows) were traced over 24.5 hours with their daughter cells indicated by the same colored arrows/arrowheads. Both sclerotome progenitors divided at least once to give rise to one perivascular fibroblast (arrowheads) as well as several interstitial cells (arrows). Snapshots of this time-lapse movie are shown in Fig 2B. *n* = 6 embryos. Scale bar: 50 μm.

**Movie S2. Perivascular fibroblasts function as pericyte progenitors.** *pdgfrb^NTR-mCherry^; col1a2:GFP* embryos were imaged from 54 hpf to 73 hpf at 6 minute intervals with time stamps indicated in the hh:mm format. Newly differentiated pericytes were retrospectively traced to identify their cell of origin. One perivascular fibroblast (green, arrows) traced can be seen gradually upregulating *pdgfrb^NTR-mCherry^* expression and extending pericyte-like cellular processes (notched arrowheads). Snapshots of this time-lapse movie are shown in Fig 4A. *n* = 7 embryos. Scale bar: 25 μm.

**Movie S3. Perivascular fibroblasts function as pericyte progenitors.** *nkx3.1^NTR-mCherry^; pdgfrb:GFP* embryos were imaged from 54 hpf to 76 hpf at 15 minute intervals with time stamps indicated in the hh:mm format. Two perivascular fibroblasts traced (arrows) can be seen gradually upregulating *pdgfrb:GFP* expression and extending pericyte-like cellular processes (notched arrowheads). Note that one of the traced cells (top arrows) migrated away from the ISV at 15:15. *n* = 16 embryos. Scale bar: 25 μm.

